# In-depth analysis reveals complex molecular etiology of idiopathic cerebral palsy

**DOI:** 10.1101/2020.08.17.255158

**Authors:** Na Li, Pei Zhou, Hongmei Tang, Lu He, Xiang Fang, Jinxiang Zhao, Xin Wang, Yifei Qi, Chuanbo Sun, Yunting Lin, Fengying Qin, Miaomiao Yang, Zhan Zhang, Caihua Liao, Shuxin Zheng, Xiaofang Peng, Ting Xue, Qianying Zhu, Yan Li, Liru Liu, Jingyu Huang, Li Liu, Changgeng Peng, Dingding Han, Dong Liu, Kaishou Xu, (Cougar) Hao Hu

## Abstract

Cerebral palsy (CP), the most prevalent physical disability in children, has long been ignored with regard to its inherent molecular mechanism. In this work, we performed in-depth clinical and molecular analysis on 120 idiopathic CP families, and identified in half of the patients the underlying risk factors. By a compilation of 117 CP-related genes, we recognized the characteristic features in terms of inheritance and function, and proposed a dichotomous classification system according to the expression patterns. In two patients with both CP and intellectual disability, we revealed that the defective TYW1, a tRNA hypermodification enzyme, caused microcephaly and problems in motion and cognition by hindering neuronal proliferation and migration. We developed an algorithm and proved in mice brain that this malfunctioning hypermodification specifically perturbed the translation of a subset of proteins involved in cell cycling. In a CP patient with normal intelligence, we identified a mitochondrial enzyme GPAM, the hypomorphic form of which led to hypomyelination of corticospinal tract. We confirmed that the aberrant Gpam in mice perturbed the lipid metabolism in astrocyte, resulting in suppressed astrocyte proliferation and a shortage of lipid contents supplied for oligodendrocyte myelination. This work broadened the scope of understanding of CP etiology and paved a way for innovative therapeutic strategies.

## Introduction

Cerebral palsy (CP) is an umbrella term spanning a group of disorders with compromised ambulant performances, which is attributed to non-progressive disturbance in the developing fetal and infant brains ^1^. CP can be categorized according to motor signs (spasticity, dyskinesia, ataxia, and hypotonia, among which spasticity manifests in ~80% of CP cases), extremities involvement (hemiplegia, monoplegia, diplegia, triplegia, and quadriplegia), and anatomical sites of the brain lesion (cerebral cortex, pyramidal tract, extrapyramidal system, and cerebellum) ^2,3^. In CP children 2 years or older, severity is reliably classified by using the 5-level Gross Motor Function Classification System (GMFCS) ^4^. The common comorbidities of CP include intellectual disability (ID), epilepsy, sensorial impairments, language deficit, autism spectrum disorders (ASD), sleep disorders, and secondary musculoskeletal defects ^2,3^. CP is the most prevalent physical disability of childhood, with a prevalence of 2-3.5 per 1,000 live births in countries with varying socioeconomic development (more than 20 million patients around the world), and the incidence of CP at term remains consistent for 50 years ^1–3,5–7^. CP poses a critical problem for rehabilitation and welfare systems, and the lifetime costs for a CP child in USA are estimated to be $1 million ^5^. The conventional risk factors of CP are believed to be gestational and perinatal insults, including birth asphyxia, neonatal infections, and teratogens ^1,5^. However, two thirds of CP patients are born at term, and birth asphyxia happens in less than 10% of CP cases ^2,6,7^. Because the causality of CP is not confirmed in 80% of cases, it is reasonable that there exists uncharted mechanism accounting for the majority of CP events ^3^.

A growing body of evidence indicates prevailing genetic contributors to the etiology of CP, including the familial cases, the high concordance rate of monozygotic twins, the association with increased parental ages, and the prevalence of congenital malformation in CP patients ^8–12^. Large cohorts of case-control studies assessing polymorphism in a variety of candidate genes did not identify any significantly associated variants ^2,13,14^. Therefore, the genetic underpinnings of CP are likely similar to that of other neurodevelopmental disorders (NDD) such as ID and ASD, where causative variants are rarely detected in association studies but with large effect size ^15–17^. In line with this hypothesis, clinically relevant copy number variation (CNV), both *de novo* and inherited, were recently identified in four collections of CP patients, resolving 4% - 31% of the cryptogenic cases ^18–21^. Meanwhile, gene-panel sequencing or whole-exome sequencing revealed pathogenic variants, mostly *de novo*, in an aggregate of five CP studies, yielding molecular diagnosis for ~16% of cases ^22–26^.

So far, the genetic and molecular information on CP etiology has been insufficient, which impedes an improvement in prophylaxis, prognosis, and treatment of CP patients in the era of precision medicine. Large cohorts need to be recruited to fully decipher the gene spectrum due to the high genetic heterogeneity of CP, and geographic and ethnic bias should be addressed, since most of the reported studies were conducted in the Caucasian populations. Whole-genome sequencing, instead of gene-panel sequencing or whole-exome sequencing, can render better coverage of predisposing alleles, and somatic mosaicism or post-zygotic mutations (PZM) in CP patients must be highlighted as in other NDDs ^27,28^. An oriented investigation into the common characteristics of CP-related genes, plus detailed mechanistic studies, seems worthwhile, before the introduction of a gene-based classification system that will have profound impact on the clinical practice.

## Results

### CP cohort recruitment, medical evaluation, and genetic screening

The 120 CP families, including 118 trios and 2 quartets, were recruited by the Department of Rehabilitation of the Guangzhou Women and Children’s Medical Center (GWCMC) from August 2014 to June 2017 in a cohort study. This study was approved by the ethic committee of GWCMC, and the written informed consents were signed by the participating families. The following inclusion criteria were adopted: (1) a diagnosis of CP made by a board of pediatric neurologists, typically based on the early-onset disabling non-progressive pyramidal and/or extrapyramidal signs; (2) an age of 2 years and older; (3) uneventful pregnancy and delivery (most of the CP patients were born at term). The following exclusion criteria were implemented: (1) perinatal insults including asphyxia, head trauma, encephalitis, and brain tumor; (2) late-onset hereditary spastic paraplegia (HSP) and DOPA-responsive dystonia (DRD), due to the distinct progression trajectory and featured fluctuation of manifestation; (3) maternal infections such as rubella and cytomegalovirus; (4) hyperbilirubinemia. The classification and summary of the CP cohort were shown in Figure 1 (A) and (B). As expected, the majority of cohort patients belonged to spastic CP, followed by dyskinetic and ataxic CP. Patients with severe motor impairment (GMFCS level IV and V) were less than those mildly disabled in motion (GMFCS level I - III). There was a significant bias for male in this CP cohort, as observed previously in ID and ASD cohorts ^29^. The prevalent comorbidities included ID/ASD and microcephaly. More details of the CP patients can be found in Supplementary table 1 and Supplementary figure 1.

**Figure 1:**
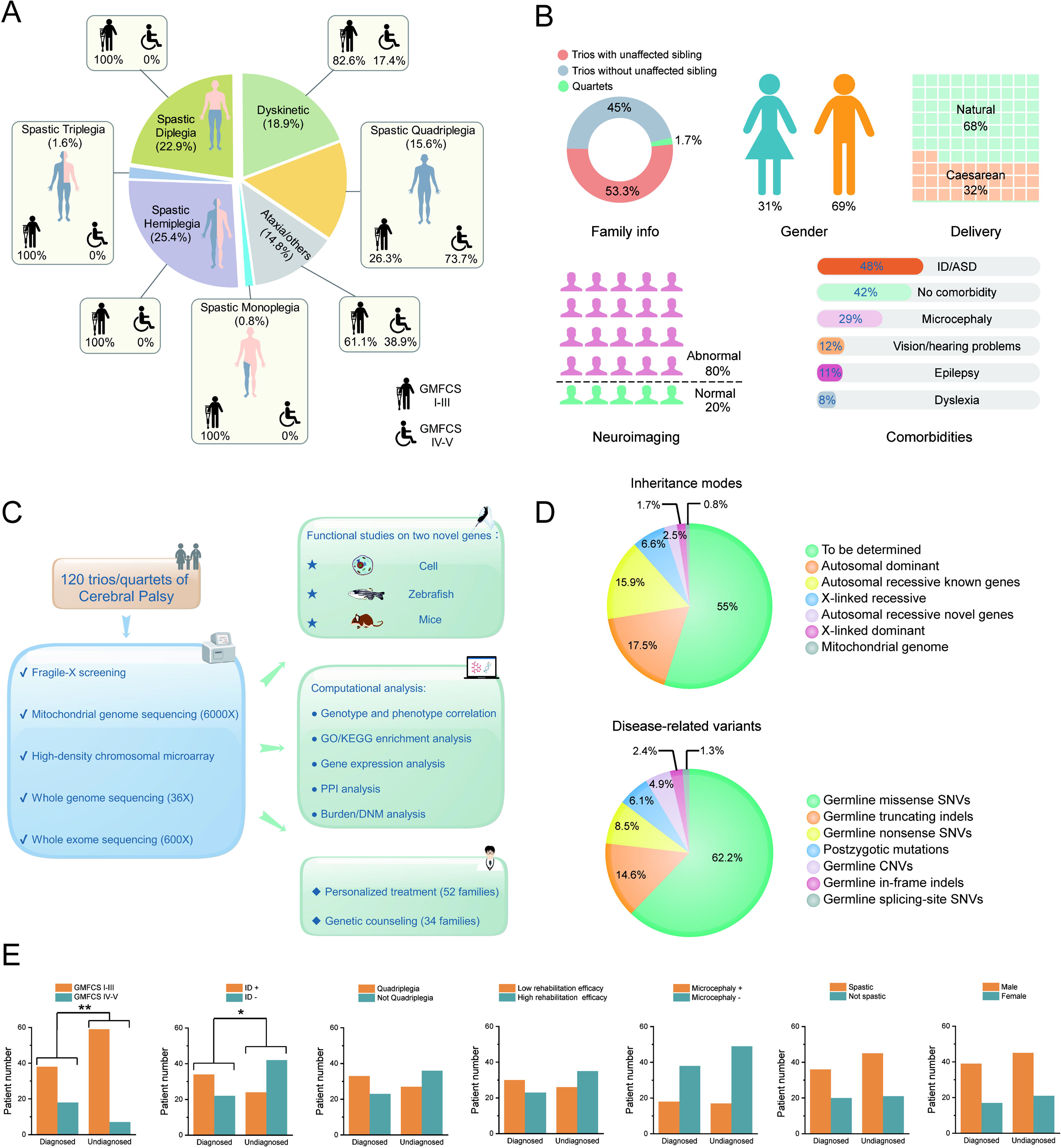
CP cohort study information. **(A)** Classification of the cohort patients based on motor signs, anatomical distribution, and motor severity. **(B)** Summary of the cohort patients in terms of family information, gender, delivery, neuroimaging, and comorbidities. **(C)** Project schematics, including cohort recruitment, genetic assessment, gene-oriented bioinformatics and functional analysis, and gene-based clinical practice. **(D)** Inheritance modes and disease-related variants identified in this study. **(E)** Correlations between diagnostic rate and clinical parameters. ID+: with intellectual disability. ID-: without intellectual disability. Microcephaly+: with microcephaly. Microcephaly-: without microcephaly.

As shown in Figure 1 (C) and Supplementary table 2, we initiated a comprehensive genetic assessment by multiple platforms, including targeted PCR for fragile-X, mitochondrial genome sequencing (6000X), high-density cytogenetic microarray, whole-exome sequencing (600X), and whole-genome sequencing (36X). In principle, we defined the disease-related variants by adopting the ACMG recommended standards, reinforced by an in-house variation database of 2,247 ethnic-matched unaffected individuals, and another database (https://db.cngb.org/cmdb/) of >50,000 individuals of East Asian descent ^30,31^.

In sum, we identified CP-related candidate genes and variants in 55 families (~45% of the 120 CP families), a comparable rate with that of other NDDs ^16,32–35^. Interestingly, the unexpected abundance of recessive variants (about a quarter) in the candidate genes, was notably higher than that of the previous NDDs investigations in outbred populations ^33,36,37^. Whether this was attributed to the specific genetic structure of ethnic group in this study, or due to the characteristic etiology of CP *per se*, remained to be solved in future study ^38^.

Of the 82 CP-related variants identified in this cohort, germline point substitution took up ~72%, followed by germline indels (~17%) and CNVs (~5%) (Figure 1 (D)). Post-zygotic mutations (PZMs) were for the first time described in a CP cohort, in which ~6% of the CP-related variants were classified as PZMs. This finding met with the expectation derived from other NDDs studies, and filled a gap in the understanding of NDDs etiology ^27^.

A closer look at the phenotype-genotype correlation in the patients revealed that: patients with more severe physical impairment (GMFCS level IV or V) had a higher chance to harbor CP-related variants (P-value=0.00613, Fisher exact test), than those patients less disabled in movement (GMFCS level I - III); patients with comorbidity of ID were more likely to have deleterious variants (P-value=0.01065, Fisher exact test), compared to those without ID (Figure 1 (E)). We also noticed the raised diagnostic rates in the quadriplegia patients and the patients with poor rehabilitation outcome, although the statistical powers were not strong enough (P-value=0.09919 and 0.1884, respectively, Fisher exact test).

In this cohort, we found altogether 129 protein-changing *de novo* mutations (DNMs) in 117 CP patients, significantly higher than the in-house 2,247 ethnic-matched unaffected individuals (P-value=0.00375, unpaired t-test). Paternal ages were revealed to correlate positively with DNM counts of the patients (r=0.63, P=0.025, Pearson correlation test). We also detected 27 PZMs in 66 CP patients, but no significant correlation with paternal or maternal ages was established. When counting only private mutations of the CP families as the data input of a burden analysis, we concluded that the 146 private mutations found in 117 families were significantly higher than what we observed in the in-house 2,247 ethnic-matched controls (P-value=0.00126, unpaired t-test).

The details of disease-related variants identified in this study are shown in Table 1 and Supplementary table 3.

**Table 1:**
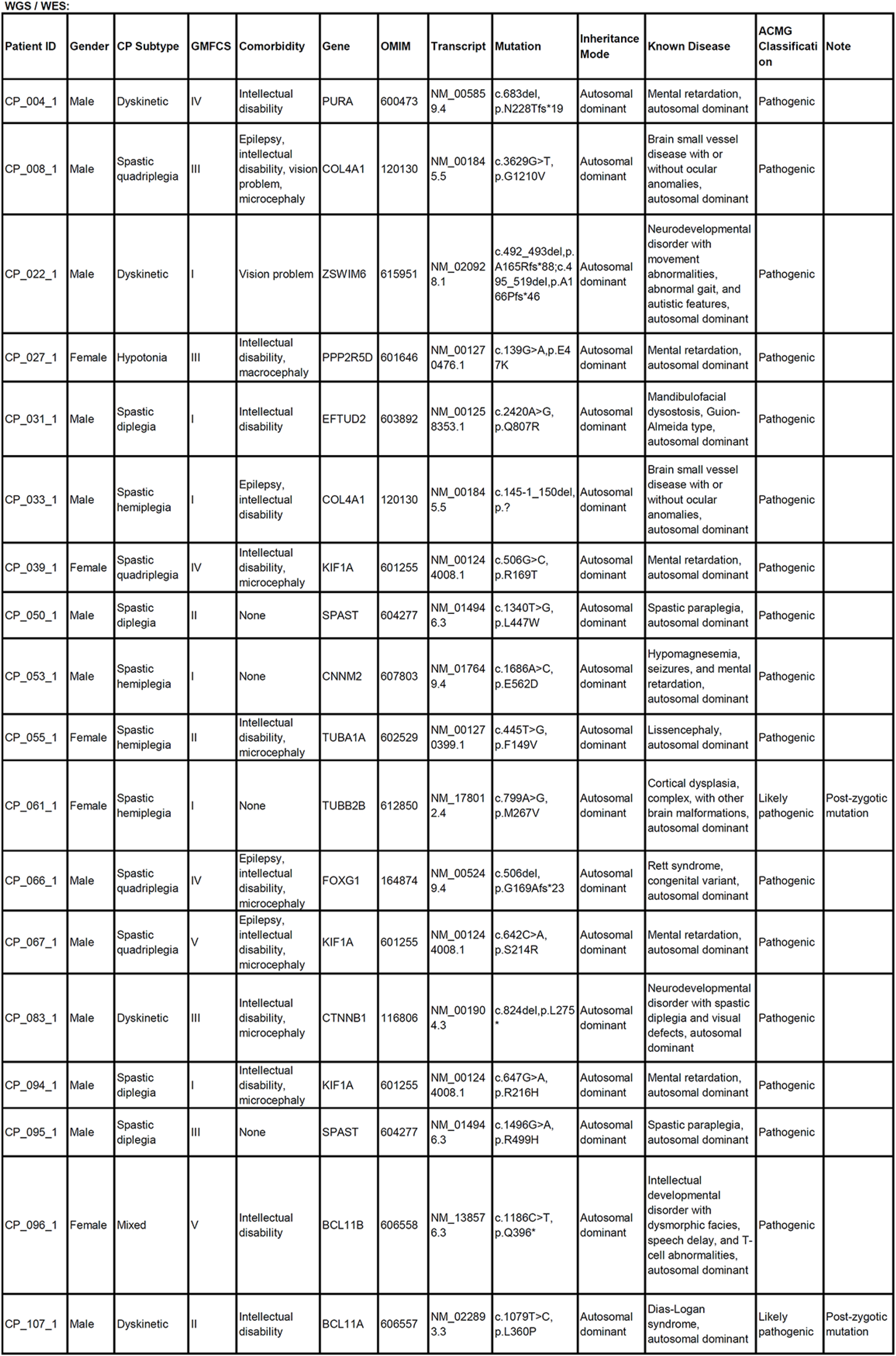
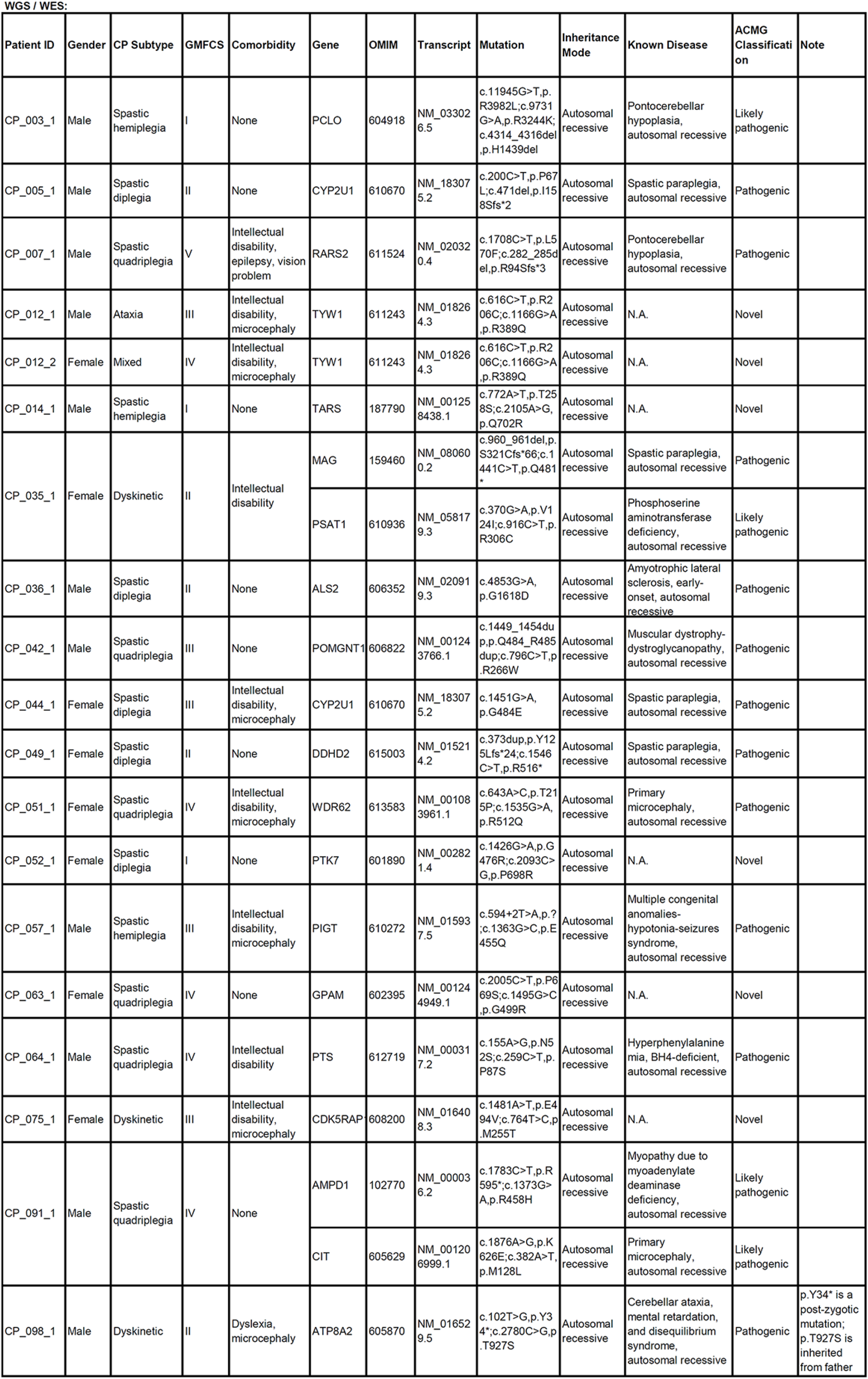
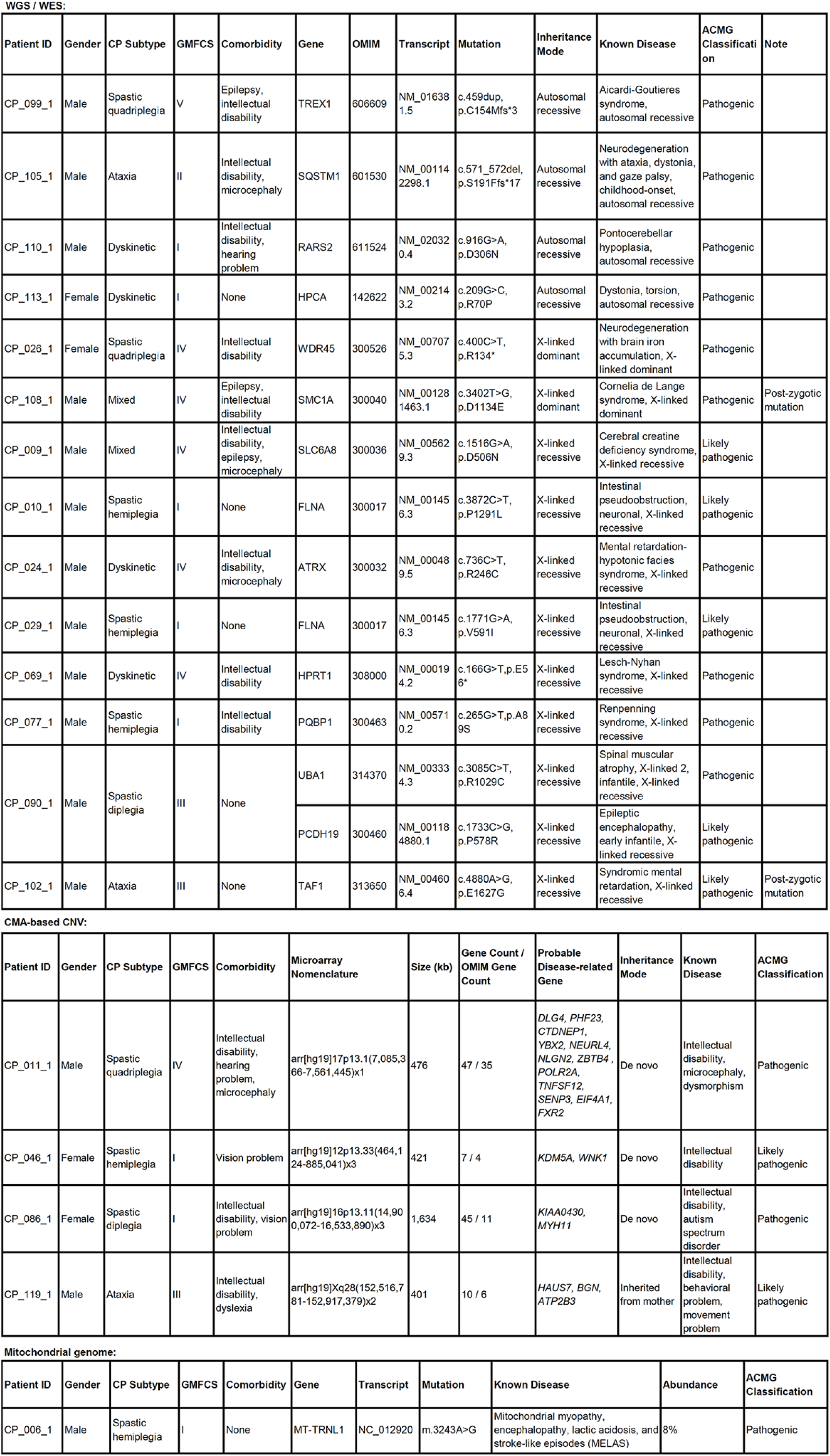
CP-related variants identified in the cohort. The variants were sorted by the platforms and inheritance modes. Motion disability was scored by the Gross Motor Function Classification System (GMFCS). Known disease for a specific gene was retrieved from the Online Mendelian Inheritance in Man (www.omim.org).

### Assessment of the role of *de novo* mutations

On aggregate, we characterized 26 CP-related *de novo* events in about one fifth of the cohort individuals, which included 18 autosomal dominant mutations (2 post-zygotic), 2 X-linked dominant mutations (1 post-zygotic), 3 *de novo* CNVs spanning multiple disease-related genes, and 1 *de novo* mutation in the mitochondrial genome (Supplementary figure 2 and 3). It is noted that there was a PZM observed on a male patient’s (CP_102_1) chromosome X while the affected gene *TAF1* (OMIM: 313650) is known to be related to X-linked recessive ID. Also interestingly, a patient (CP_098_1) had compound heterozygous variants in the gene of *ATP8A2* (OMIM: 605870), albeit with one allele of post-zygotic origin.

We found in two spastic diplegia patients (CP_050_1 and CP_095_1) DNMs of *SPAST* (OMIM: 604277), a gene frequently identified in CP patients that is related to early-onset autosomal dominant spastic paraplegia. We also identified in three CP patients (CP_039_1, CP_067_1, CP_094_1) DNMs of *KIF1A* (OMIM: 601255), which is known to cause autosomal dominant mental retardation and motor delay. These patients showed variable phenotype (spastic diplegia and quadriplegia, GMFCS level I - V). Nevertheless, all of the three patients manifested ID and severe microcephaly. Another recurrent gene in this cohort is *COL4A1* (OMIM: 120130), in which we identified DNMs in two CP patients (CP_008_1 and CP_033_1). Both patients presented spastic CP, epilepsy, and severe ID. In addition, one patient (CP_008_1) showed severe microcephaly and vision problem.

Some of the CP patients harbored disease-related variants in genes that are functionally relevant. For example, the patient CP_107_1 (dyskinetic CP with ID) had a post-zygotic *de novo* mutation of *BCL11A* (OMIM: 606557), while the patient CP_096_1 (mixed-type CP with ID) had a *de novo* truncating mutation of *BCL11B* (OMIM: 606558). *BCL11A* and *BCL11B* encode zinc-finger proteins that regulate transcription, highly expressed in brain and bone marrow, and were reported to be related to autosomal dominant developmental disorders. Similar scenario occurred in two CP patients (CP_055_1 and CP_061_1), where the former (spastic hemiplegia with severe ID and microcephaly) had a *de novo* missense mutation of *TUBA1A* (OMIM: 602529), and the latter (spastic hemiplegia with mild motor impairment) had a post-zygotic *de novo* mutation of *TUBB2B* (OMIM: 612850). *TUBA1A* and *TUBB2B* code for brain-enriched tubulin components as subunits of microtubules, and were known to be associated with autosomal dominant brain malformations.

### Identification of the variants involved in recessive disease

Altogether we confirmed recessive-disease-related variants in 32 CP families (one quarter of the 120 cohort families), including 22 families with autosomal recessive disease, 9 families with X-linked recessive disease, and 1 CNV located at Xq28 of a male patient (CP_119_1) inherited from the unaffected mother, where a variety of CNVs were reported previously in patients of neuropsychiatric disorders, in some cases, with gender bias ^39,40^.

We noticed that, among the 22 CP families with autosomal recessive disease, the zygotic ratio of compound heterozygosity vs. homozygosity was 3:1, which was acceptable in a largely outbred country. Interestingly, among the 6 homozygous variants, only one was proved to be autozygous by the CMA-based genotyping and family history consulting (CP_036_1, second-cousin consanguinity). This observation indicated the existence of CP-related hotspot variants in a specific population.

The defects of *RARS2* (OMIM: 611524), a gene coding for mitochondrial arginyl-tRNA synthetase, led to manifestation in two patients (CP_007_1 and CP_110_1). The former patient had severe spastic quadriplegia with ID, epilepsy, and vision problem, probably caused by the highly pathogenic compound heterozygous variants of *RARS2*; whereas the latter patient, with mild dyskinetic CP, moderate ID and hearing problem, was likely associated with a homozygous missense variant that may be less damaging.

The patient CP_105_1 presented ataxic CP with mild ID and microcephaly. A homozygous truncating variant was found in the gene of *SQSTM1* (OMIM: 601530), encoding a ubiquitin-binding protein. According to a recent report, four families of autosomal recessive childhood-onset neurodegenerative disorder were found to have detrimental variants in *SQSTM1* ^41^. However, the onset age in our case was much younger (in the first year since birth), therefore, this finding expanded the phenotype spectrum related to the gene of *SQSTM1*.

*TARS* (OMIM: 187790) codes for a threonyl-tRNA synthetase and plays dual roles both as an assembly scaffold of translation initiation components and as a target mRNA selector ^42^. A very recent report introduced two individuals carrying *TARS* variants with trichothiodystrophy ^43^, while we found a spastic hemiplagia patient (CP_014_1) with biallelic *TARS* variants, who presented overlapping phenotype including developmental delay, but without manifestation in hair or skin (Supplementary figure 4).

*PTK7* (OMIM: 601890) encodes a tyrosine kinase of Wnt-signaling pathway, and altered PTK7 activity in mice models caused perturbation of neural tube development ^44^. Very recent studies found association of *PTK7* variants with neural tube defects and fetal anomaly in human ^45,46^. We found that the patient CP_052_1 had compound heterozygous deleterious variants of this gene, and the patient showed spastic diplegia with mild motion impairment (Supplementary figure 5).

*CDK5RAP1* (OMIM: 608200) encodes an enzyme involved in the mitochondrial tRNAs modification resulting in stabilization of codon/anticodon interaction, and is primarily expressed in neurons of the central nervous system. Defects of this gene were reported to cause myopathy in mice and humans ^47^. Biallelic damaging variants of this gene were revealed in CP_075_1, a dyskinetic CP patient with ID and microcephaly (Supplementary figure 6).

Another two novel candidate genes, namely, *TYW1* (OMIM: 611243) and *GPAM* (OMIM: 602395), were identified in two CP families with distinct phenotypes (CP_012 and CP_063, respectively). The former family had multiple patients, manifesting both CP and ID; while the latter family had a patient with CP but without ID. Because this dichotomous manifestation epitomized the two major subgroups of CP (with or without ID), we carried out detailed functional studies, and presented the results in the subsequent parts of this manuscript.

### Common features of CP-related genes

When lumping CP-related genes from this cohort and from literature, there have been in sum 117 CP-related genes identified up till now (Supplementary table 4). This number is far less than the 700+ genes involved in ID and other NDDs ^29^. About half of the CP-related genes are defined to be ID/NDD genes, and this is not unexpected since CP is on a diagnostic continuum with ID/NDD (Figure 2 (A)). The inheritance modes of the 117 CP-related genes include autosomal dominant (42.7%), autosomal recessive (42.7%), X-linked recessive (12%), and X-linked dominant (2.6%). Compared to ID/NDD genes, CP-related genes lack significantly in autosomal recessive genes but abound in autosomal dominant genes (P-value=0.0006, Fisher exact test) (Figure 2 (A)). We thus speculated that there was a great chance for novel autosomal recessive CP-related genes to be identified in future studies.

**Figure 2:**
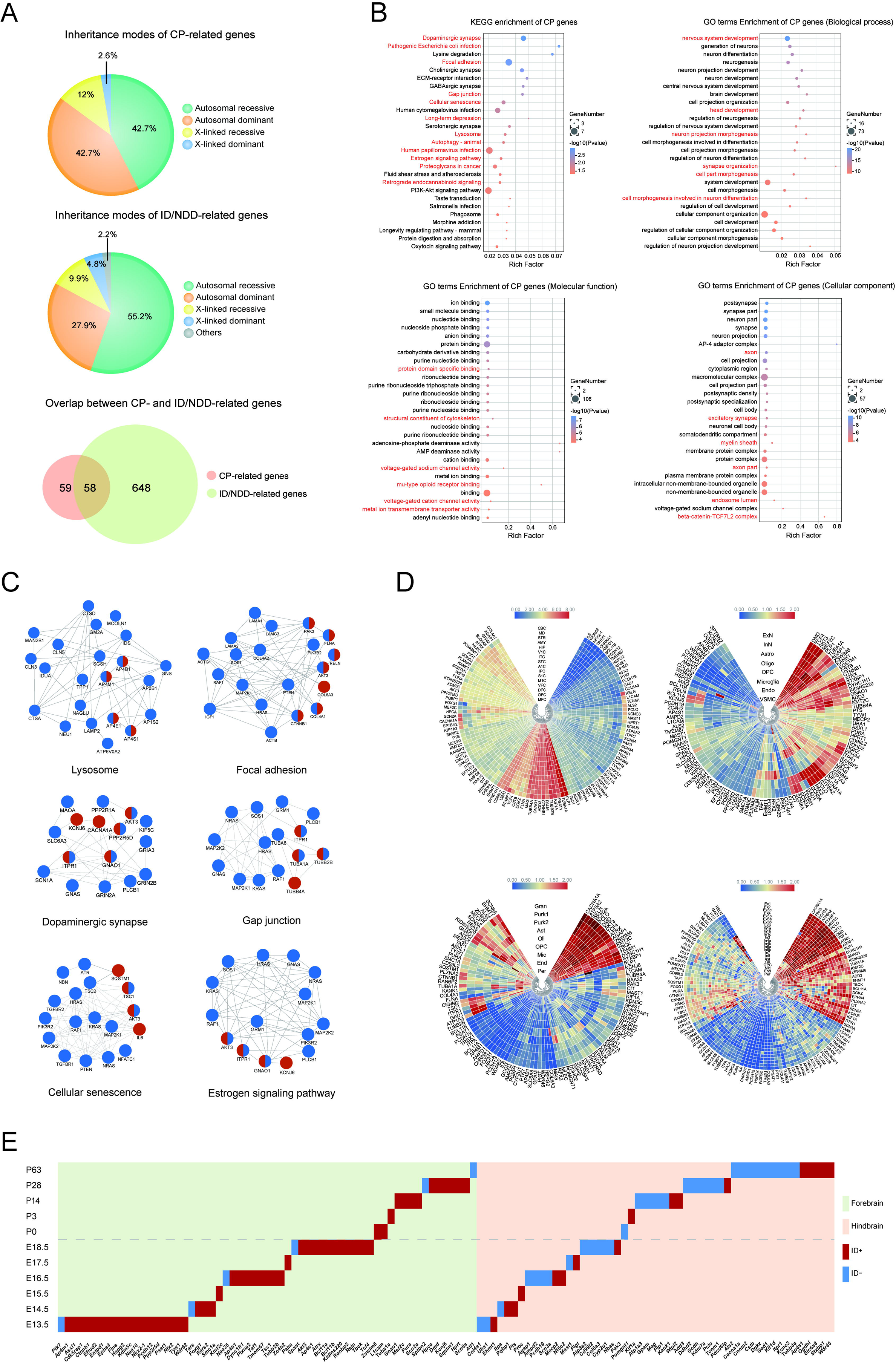
CP-related gene summary. **(A)** Inheritance modes comparison between the CP-related genes and ID/NDD-related genes. The data of CP-related genes were from this study and a collection of literature. The data of ID/NDD-related genes were retrieved from literature ^29^. **(B)** Enriched KEGG pathways and GO terms of the CP-related genes. Upper left: enriched KEGG pathways. Upper right: enriched GO terms of Biological Process. Bottom left: enriched GO terms of Molecular Function. Bottom right: enriched GO terms of Cellular Component. Red color indicated the pathways and GO terms also enriched in the ID/NDD-related genes. **(C)** Protein-protein interaction network modules linking CP-related genes (red), ID/NDD-related genes (blue), and shared genes (half red, half blue). **(D)** Expression of CP-related genes in different human brain regions and cell types. Upper left: expression heatmap in 16 human brain regions ^50^. MFC: medial prefrontal cortex. OFC: orbital prefrontal cortex. DFC: dorsolateral prefrontal cortex. VFC: ventrolateral prefrontal cortex. M1C: primary motor cortex. S1C: somatosensory cortex. IPC: posterior inferior parietal cortex. A1C: primary auditory cortex. STC: superior temporal cortex. ITC: inferior temporal cortex. V1C: primary visual cortex. HIP: hippocampus. AMY: amygdala. STR: striatum. MD: mediodorsal nucleus of thalamus. CBC: cerebellar cortex. Upper right: expression heatmap in 8 cell types of human dorsolateral prefrontal cortex ^50^. ExN: excitatory neuron. InN: interneuron. Astro: astrocyte. Oligo: oligodendrocyte. OPC: oligodendrocyte progenitor cell. Endo: endothelial cell. VSMC: vascular smooth muscle cell. Bottom left: expression heatmap in 9 cell types of human cerebellum ^51^. Gran: cerebellar granule cell. Purk1: Purkinje neuron, subtype 1. Purk2: Purkinje neuron, subtype 2. Ast: astrocyte. Oli: oligodendrocyte. OPC: oligodendrocyte precursor cell. Mic: microglia. End: endothelial cell. Per: pericyte. Bottom right: expression heatmap in 24 cell types of human frontal cortex ^51^. Ex1: excitatory neuron, subtype 1. Ex2: excitatory neuron, subtype 2. Ex3e: excitatory neuron, subtype 3e. Ex4: excitatory neuron, subtype 4. Ex5b: excitatory neuron, subtype 5b. Ex6a: excitatory neuron, subtype 6a. Ex6b: excitatory neuron, subtype 6b. Ex8: excitatory neuron, subtype 8. In1a: inhibitory neuron, subtype 1a. In1b: inhibitory neuron, subtype 1b. In1c: inhibitory neuron, subtype 1c. In3: inhibitory neuron, subtype 3. In4a: inhibitory neuron, subtype 4a. In4b: inhibitory neuron, subtype 4b. In6a: inhibitory neuron, subtype 6a. In6b: inhibitory neuron, subtype 6b. In7: inhibitory neuron, subtype 7. In8: inhibitory neuron, subtype 8. Ast: astrocyte. Oli: oligodendrocyte. OPC: oligodendrocyte precursor cell. Mic: microglia. End: endothelial cell. Per: pericyte. **(E)** Expression climax of CP-related genes in mice, with regard to brain regions (forebrain and hindbrain), developmental stages (11 time points before and after birth), and comorbidity (with or without ID). X-coordinate: CP-related genes; Y-coordinate: embryonic and postnatal days. Gene expression data were retrieved from https://www.ebi.ac.uk/gxa/home.

We found that the 117 CP-related genes were enriched in a variety of pathways and protein-protein interaction modules, involving multiple aspects of brain development and functioning (Figure 2 (B). We noticed that there were at least 6 functional modules that reflected the essential and coordinated roles of CP-related genes in living organisms, i.e., lysosome, gap junction, dopaminergic synapse, focal adhesion, cellular senescence, and estrogen signaling modules (Figure 2 (C)). Estrogen signaling pathway regulates a plethora of physiological processes in mammals, including cellular homeostasis and behavior, given the significant gender bias in populations of CP, ID and ASD, to explore its involvement in the etiology of NDDs would be an interesting topic. On the other hand, we noticed the pathogen-related modules (pathogenic *escherichia coli* infection, human cytomegalovirus infection, and human papillomavirus infection) enriched in the CP/ID/NDD genes, comprising ~11% of the 117 CP-related genes, and occurring in the CP cohort patients (4%), such as *TUBA1A*, *TUBB2B*, *CTNNB1*, *COL4A1*. Although it is well known that CP and ID can be sequelae of intracranial pathogen infection, whether the variants of these genes increase the relevant disease risk remains elusive.

When data-mining in the public databases and literature of gene expression profiles in brain, we were aware that the CP-related genes were expressed in a variety of anatomic structures, including prefrontal cortex, motor cortex, cerebellar cortex, hippocampus, striatum, somatosensory cortex, etc, and in a variety of brain cells, including excitatory neuron, inhibitory neuron, interneuron, astrocyte, oligodendrocyte, cerebellar granule cell, Purkinje neuron, microglia, etc (Figure 2 (D)). The top3 brain regions were cerebellum, primary motor cortex, and striatum; the top3 cell types were Purkinje cell, astrocyte, and oligodendrocyte. For example, *TYW1* showed enriched expression in neuronal cells in each brain region, but not observable in pericytes or microglia cells. *GPAM* showed exclusively high expression in astrocytes, whereas untraceable in other brain cells. Expression of *CDK5RAP1* was observed in several subtypes of inhibitory neuron, vascular smooth muscle cells and endothelial cells, the latter two reflecting relationship with brain angiogenesis. *PTK7* showed higher expression in pericytes compared to other cell types, indicating a role in capillary formation. The specific expression patterns can serve as useful guidelines for further mechanistic studies on these genes.

As it was found, CP-related genes reached highest expression in different brain regions (forebrain 57%, hindbrain 43%) and at sequential developmental stages (embryonic 58%, postnatal 42%) (Figure 2 (E)). Interestingly, we noticed that CP-related genes could be roughly divided into two subgroups with distinctive features, namely, forebrain-expressed genes and hindbrain-expressed genes. The former genes, such as *TYW1*, highly expressed in forebrain including motor cortex and striatum, had a significantly higher chance to have comorbidity of ID than the latter ones, such as *GPAM*, which were specifically expressed in hindbrain including brainstem and cerebellum (P-value=0.002, Fisher exact test). A spin-off observation was that, forebrain-expressed CP-related genes preferred to reach expression height during embryonic periods, while hindbrain-expressed CP-related genes approached expression climax throughout embryonic and postnatal periods, but with an inclination for postnatal days (P-value=0.05, Fisher exact test). It is an intuitive surmise that, the forebrain-expressed genes should participate in neurogenesis that is generally finished before birth, while the hindbrain-expressed genes may take part in myelination that is not complete until two years after birth in human ^48,49^.

Next, we will illustrate this dichotomous classification system by delving into two typical CP-related genes, newly identified in this study, namely, *TYW1* and *GPAM*.

### Null and hypomorphic alleles of *TYW1* caused primary microcephaly and impairment in motion and cognition

The male proband CP_012_1, an ataxic CP patient, was the second of two children of a non-consanguineous, healthy Chinese couple with unremarkable family history. He was born by normal delivery at 40 weeks of gestational age after an uneventful pregnancy with a birth weight of 3.8 kg (+1.17 S.D.). At conception, the mother was aged 31 and the father was 34 years old. By clinical examination at age of 6, the patient’s height, body weight, and head circumference were 103 cm (−3.20 S.D.), 16 kg (−2.06 S.D.), and 47.5 cm (−3.26 S.D.), respectively. At age of 8 years 7 months, the cranial MRI of the patient showed a slightly widened cerebral subarachnoid space, enlarged ventricle, and abnormal cerebellum morphology. Cognitive evaluation (Wechsler Preschool and Primary Scale of Intelligence) was performed at age of 7 years and 6 months, with full-scale IQ 40, indicating a moderate ID. The level of Gross Motor Function Classification System (GMFCS) was III, i.e., walk using a hand-held mobility device. The fine motor skill was poor since the Manual Ability Classification System (MACS) was evaluated at the level of IV (handles a limited selection of easily managed objects in adapted situations). Another patient CP_012_2, a three-year-older sister of the proband, was diagnosed mixed-type CP. The mother had a normal delivery after 40 weeks of uneventful gestation, and the birth weight of the patient was 3.2 kg (−0.02 SD). The parameters of clinical examination at age of 14 years of the patient were: height 150 cm (−1.51 S.D.), weight 41 kg (−1.03 S.D.), and head circumference 49 cm (−3.64 S.D.). Cognitive evaluation (Estimated Cognitive Level of Children with CP) was performed at age of 13 years and 1 month with IQ <50, indicating a moderate ID. The level of the GMFCS was IV, meaning self-mobility with limitations and needing powered mobility. The fine motor skill was poor since the MACS was at level III (handles objects with difficulty, need help to prepare and/or modify activities). The medical information of the two patients was presented in Figure 3 (A) and Supplementary table 1 and 5.

**Figure 3:**
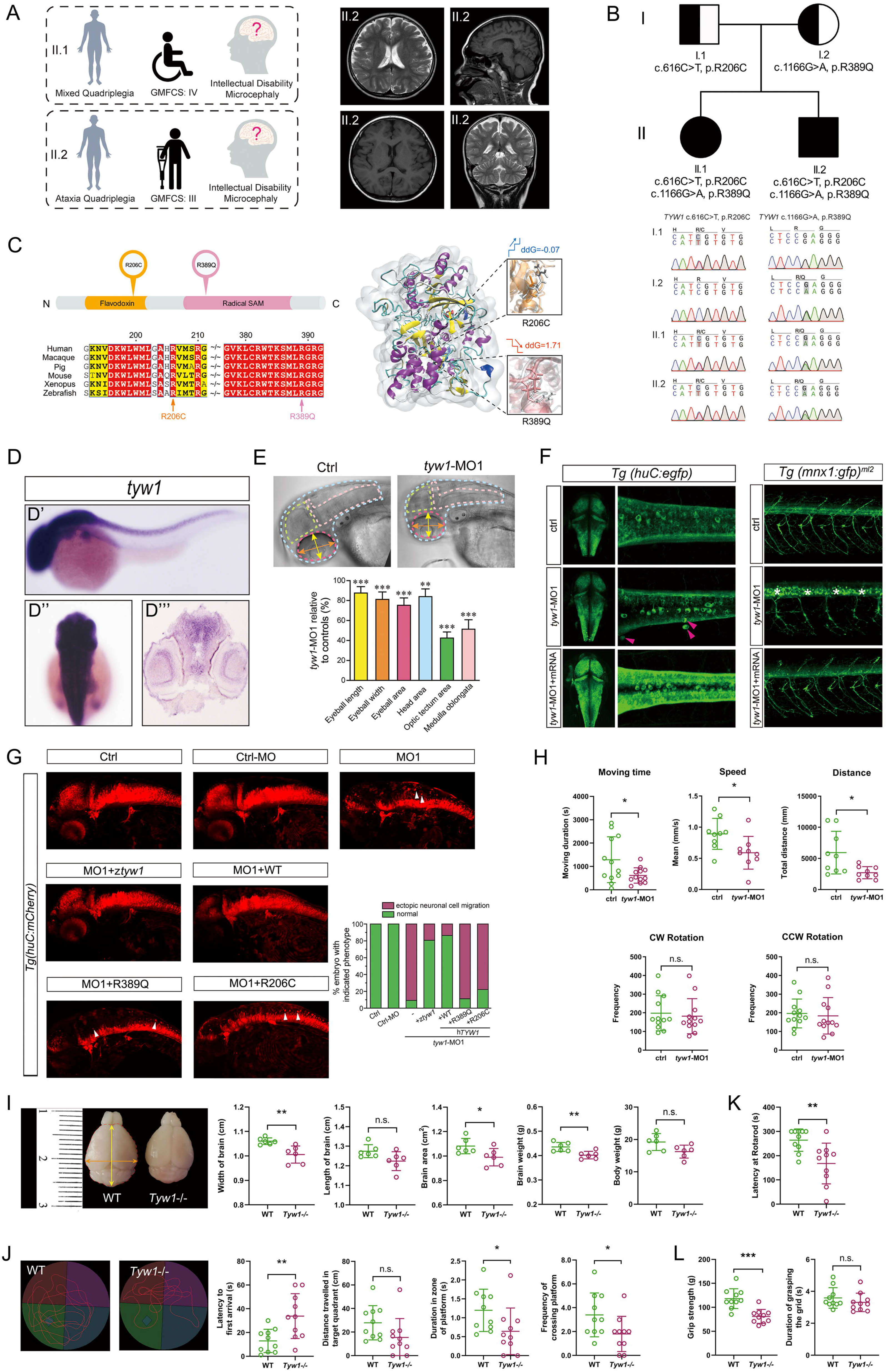
Correlation between phenotype and genotype of *TYW1*. **(A)** Clinical summary of the two index patients harboring *TYW1* variants, and the cranial magnetic resonance imaging of the patient II.2 (CP_012_1). **(B)** Compound heterozygous variants of *TYW1* were identified in the two index patients, which co-segregated with the phenotype in the family members. **(C)** The two identified amino-acid changes, i.e., R206C and R389Q, were located inside the protein domains of flavodoxin and radical SAM, respectively. The two changes took over the highly conserved amnio-acid positions throughout a series of species. Three-dimensional structural model of TYW1 protein displayed the stability alteration (ddG, Gibbs free energy) and local structural changes, before and after the introduction of two amino-acid changes. In the two zoom-in views, white bars stood for the wild-type amino acids and color bars for the mutated ones. **(D - H)** Morphological and behavioral assays on zebrafish models. **(D)** Whole mount *in situ* hybridization of zebrafish embryos at 36 hpf using zebrafish antisense probe of *tyw1* showed specific expression of *tyw1* in the central nervous system including brain and spinal cord. D’: lateral view. D”: dorsal view. D”’: cross section. hpf: hours post fertilization. **(E)** Head area, optic tectum area, medulla oblongata area, eyeball area, eyeball length, and eyeball width of the *tyw1* knockdown (*tyw1*-MO1, a splice-blocking morpholino) zebrafish larvae at 5 dpf were significantly decreased compared with the controls. N = 10 for each test group. ** p < 0.01, *** p < 0.001, unpaired t-test. dpf: days post fertilization. **(F)** Confocal imaging of egfp positive cells in the transgenic zebrafish *Tg(huC:egfp)* embryos at 48 hpf showed ectopic neuronal cells (indicated by red arrowheads) in the *tyw1*-MO1 zebrafish group, which could be rescued by the wild-type *tyw1* mRNA. Confocal imaging of gfp positive cells in the transgenic zebrafish *Tg(mnx1:gfp)^ml2^* embryos at 72 hpf showed undifferentiated motor neuronal cells (indicated by white stars) in the *tyw1*-MO1 zebrafish group, which could be rescued by the wild-type *tyw1* mRNA. **(G)** The *tyw1* knockdown (*tyw1*-MO1) brought about ectopic neuronal cells in the zebrafish *Tg(huC:mCherry)* embryos at 48 hpf, and there were only 9% of subjects that remained normal. This abnormal phenotype could be rescued by zebrafish *tyw1* mRNA (*ztyw1*), human *TYW1* mRNA (WT), mutated human *TYW1* leading to R389Q, and mutated human *TYW1* leading to R206C, to variable extents (81%, 86%, 11%, 22%, respectively). Ectopic neuronal cells were indicated by white arrowheads. Fish number ranged from 63 to 88 in different test groups. **(H)** Swimming behavior tests of the *tyw1*-MO1 zebrafish larvae at 5 dpf compared to the controls, on moving time, speed, distance, clockwise (CW) rotation, and counterclockwise (CCW) rotation. The former three parameters revealed significant reduction of the *tyw1*-MO1 group, while the latter two parameters showed no significant change. N = 9 to 12 for each test group. * p < 0.05, n.s.: no significant difference, unpaired t-test. **(I - L)** Morphological and behavioral assays on *Tyw1* knockout (*Tyw1*−/−) mice models. **(I)** The *Tyw1*−/− mice at postnatal 8 weeks showed significantly reduced brain size and weight compared to the wild-type (WT) ones. The orange arrow line, yellow arrow line, and red dash line indicated the width, length, and area of brain, respectively. In parallel, measurement of body weight revealed no significant difference between the *Tyw1*−/− mice and the WT ones. N = 6 per genotype with equal numbers of male and female mice. * p < 0.05, ** p < 0.01, n.s.: no significant difference, unpaired t-test. **(J)** Track plots depicted the trace of mice during the probe test in a Morris water maze. The *Tyw1*−/− mice showed worse performance compared to the WT ones in the test. N = 10 per genotype with equal numbers of male and female mice. * p < 0.05, ** p < 0.01, n.s.: no significant difference, unpaired t-test. **(K)** Rotarod tests showed significantly reduced motor coordination and balance of the *Tyw1*−/− mice compared to the WT ones. N = 10 per genotype with equal numbers of male and female mice. ** p < 0.01, unpaired t-test. **(L)** Grip strength tests revealed significantly weaker grip strength of the *Tyw1*−/− mice compared with the WT mice. N = 10 per genotype with equal numbers of male and female mice. *** p < 0.001, n.s.: no significant difference, unpaired t-test.

In both of the two patients, we identified compound heterozygous variants in the gene of *TYW1* (OMIM: 611243), which were co-segregating with the phenotype in the family and confirmed by targeted PCR and Sanger sequencing (Figure 3 (B)). The pRecessive value of *TYW1* (http://exac.broadinstitute.org/) is 0.9945, a very strong indicator of recessive-disease-related gene. The two variants (NM_018264: c.616C>T, p.R206C; c.1166G>A, p.R389Q) were absent from an in-house database with 2,247 healthy controls, and were very rare in the public databases (http://exac.broadinstitute.org/; https://db.cngb.org/cmdb/) with allele frequencies below 1 × 10^−5^. Pathogenicity prediction classified the two variants as damaging or disease-causing (Supplementary table 3). Multiple protein sequences alignment among different species showed high conservation of both positions, i.e., R206 and R389, which were located in the functional domains (Figure 3 (C)). The protein structural modeling revealed the functional locations of both amino acids, and showed the reduced protein stability (R389Q) and disturbed substrate binding (R206C) (Figure 3 (C)). Compatible with the prediction, we found a dramatically lower protein level of TYW1 in the patient’s peripheral blood sample, compared to those of the parents and healthy children with same age and gender (Supplementary figure 7).

Using whole-mount *in situ* hybridization (WISH) of zebrafish, we found that *tyw1* expression was highly enriched in the developing central nervous system including brain and spinal cord (Figure 3 (D)). The morpholino-mediated *tyw1* knockdown (*tyw1*-MO) zebrafish models were established, by blocking mRNA splicing (MO1) and protein translation (MO2) of *tyw1* (Supplementary figure 8, 9 and 10). We observed significant head size reduction in the *tyw1*-MO zebrafish compared to the wild-type ones (Figure 3 (E)). It was revealed that *tyw1* deficiency in zebrafish resulted in ectopic neuronal cell migration in brain and undifferentiated motor neuronal cells in spinal cord (Figure 3 (F) and Supplementary figure 10). The phenotypic anomaly could be effectively rescued by zebrafish *tyw1* mRNA or human *TYW1* mRNA, but not by the *TYW1* mRNAs with variants identified in the CP patients (Figure 3 (G)). This result substantiated the pathogenicity of the two variants identified in the patients. By performing swimming behavior tests, we noticed that the swimming capacity of *tyw1*-MO zebrafish larvae was compromised significantly, specifically in swimming speed, time, and distance, but not in rotation (Figure 3 (H)).

By using CRISPR/Cas9 genome editing technology, we constructed a *Tyw1*-knockout mice model (Supplementary figure 11). In the *Tyw1*−/− mice, the brain size and weight were significantly below the level of those of the wild-type littermate, with abnormal intracranial morphology (Figure 3 (I) and Supplementary figure 12). Behavioral tests, i.e., the Morris water maze test, the rotarod test, and the grip strength test, all witnessed significantly reduced performances of the *Tyw1*−/− group compared to the wild-type controls (Figure 3 (J), (K), (L)). The deterioration of both motion and cognition was compatible with the manifestation of the CP patients with hypomorphic *TYW1* alleles, and reflected the underlying cerebral regions responsible for both functioning.

### Defective Tyw1 hindered neuronal proliferation and migration due to increased ribosomal frameshift in a subset of cell-cycling-related proteins

The gene of *TYW1* (OMIM: 611243) encodes a tRNA-wybutosine (tRNA-yW) synthesizing protein, localized on the cytosolic surface of the endoplasmic reticulum and responsible for producing a hypermodified guanosine, i.e., wybutosine, at the position 37 of phenylalanine tRNA (tRNA^Phe^) adjacent to the 3-prime of anticodon in archaea and eukaryotes ^52,53^. This hypermodification at postion 37 of tRNA^Phe^ is known to play a critical role in stabilizing the appropriate interaction between codon and anticodon during protein translation, absence of which incurs promoted ribosomal frameshift thus reducing translational accuracy ^54–57^. Interestingly, although there exist two codons for phenylalanine, i.e., UUU and UUC, it seems that wybutosine exerts influence only upon the codon UUU ^58^.

The expression of TYW1 are enriched in the neuronal cells, especially during the embryonic development (Figure 2 (D) and (E); Figure 4 (A) and (B)). We observed significantly decreased neurogenesis in the E13.5 mice brain when comparing the *Tyw1*−/− mice with the wild-type ones (Figure 4 (C)). In the *Tyw1*−/− mice brain, the migration of cortical neuronal precursors between E15.5 and E18.5 was hindered, resulting in obvious reduction of superficial-layer neurons and noticeable accumulation of deep-layer neurons (Figure 4 (D)). Consequently, in both frontal cortex and motor cortex of the adult mice brain, the thickness of superficial layer was significantly less in the *Tyw1*−/− group compared with the wild-type one, while the trend seemed reverse in the deep layer although the statistical power had not been gained (Figure 4 (E)). The simultaneous abnormalities in both frontal and motor cortices in the mice models provided a clue to explain the CP-plus-ID phenotype of patients with hypomorphic *TYW1* alleles in this study.

**Figure 4:**
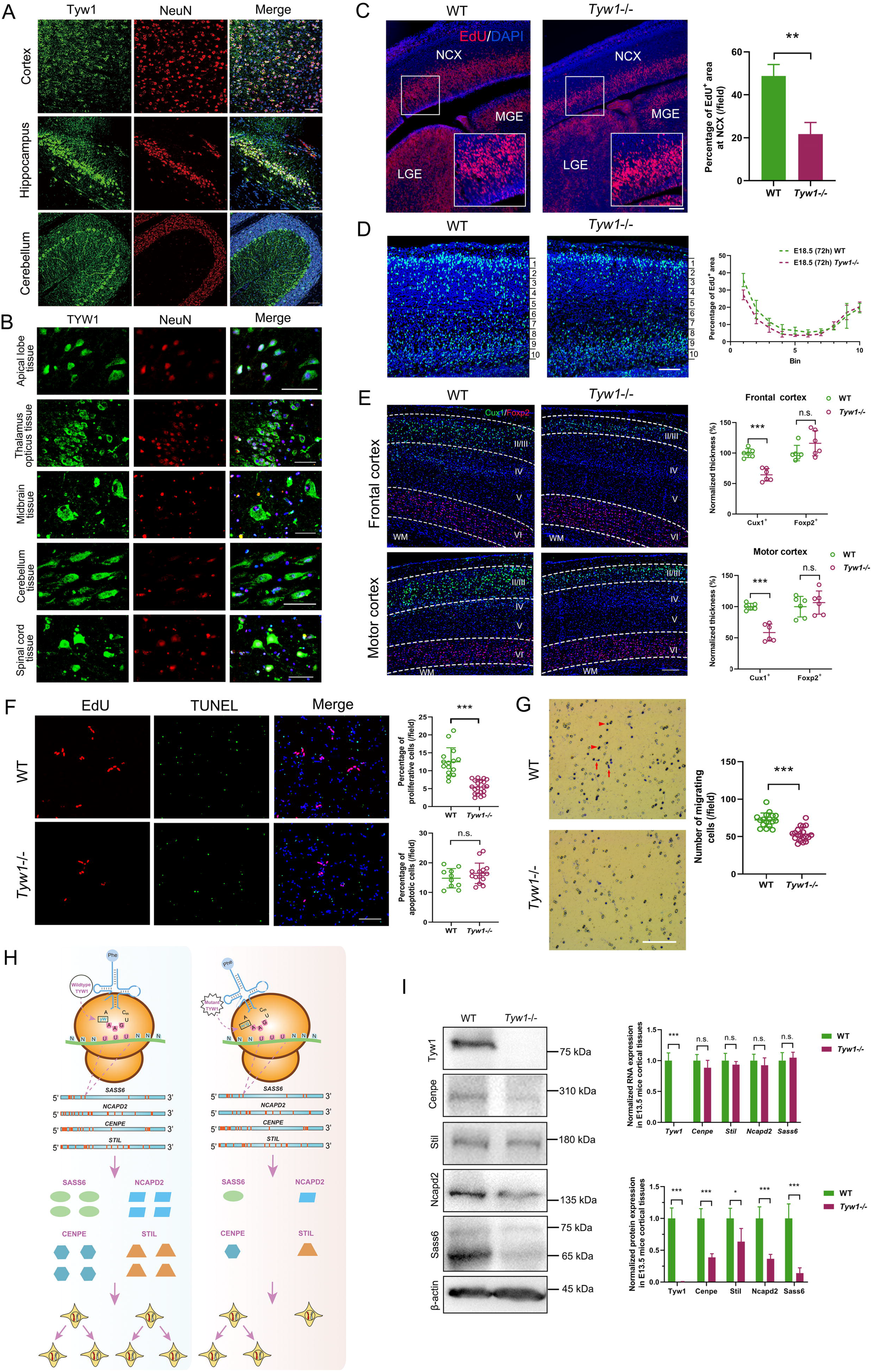
Functional mechanism of defective TYW1. **(A)** Brain section immunostaining of the wild-type mice at postnatal 8 weeks, where Tyw1 (green) and NeuN (red) were co-labeled, showed the Tyw1 expression in the neurons of cortex, hippocampus, and cerebellum. Blue: DAPI. Bar = 75 μm. **(B)** Immunostaining of the normal human brain tissue microarrays, where TYW1 (green) and NeuN (red) were co-labeled, revealed the TYW1 expression in the neurons of a variety of brain regions. Blue: DAPI. Bar = 50 μm. **(C)** EdU was intraperitoneally injected 2h before sacrifice into pregnant mice with embryos on E13.5. Cells incorporated with EdU were determined in the embryonic brain sections under fluorescence microscope and analyzed with ImageJ software. Compared to the wild-type (WT) littermate, significantly reduced percentages of EdU^+^ area were observed at neocortex (NCX) of the *Tyw1*−/− embryo brains, and at medial ganglionic eminence (MGE) and lateral ganglionic eminence (LGE) as well. Red: EdU. Blue: DAPI. N = 3 per genotype. For a specific sample, each measurement was performed twice in separate fields. ** p < 0.01, unpaired t-test. Bar = 100 μm. **(D)** EdU was injected into the pregnant mice carrying embryos on E15.5, which were observed after 72 h. The embryonic cortex was divided into ten equally spaced bins along the vertical axis. Percentage of EdU^+^ area in each bin was measured. At the *Tyw1*−/− mice cortex, EdU^+^ cells in the superficial layer were much less than that in the WT mice, while EdU^+^ cells in the deep layer were a bit more than that in the WT mice. Light blue: EdU. Blue: DAPI. N = 3 per genotype. Bar = 100 μm. **(E)** Brain section immunostaining of mice at postnatal 8 weeks, where the Cux1^+^ and Foxp2^+^ cells were labeled for neurons in superficial layer (layer II-III) and deep layer (layer VI), respectively. Compared with the WT mice, the *Tyw1*−/− mice brain showed significantly reduced thickness of the superficial layers at both frontal and motor cortices, while in the deep layers, the reverse happened, albeit without statistical significance. WM: white matter. Green: Cux1. Red: Foxp2. Blue: DAPI. N = 3 per genotype. For a specific sample, each measurement was performed twice in separate fields. *** p < 0.001, n.s.: no significant difference, unpaired t-test. Bar = 200 μm. **(F - G)** Proliferation, apoptosis and migration analysis on primary neurons from the cortex of E13.5 mice. **(F)** Quantification of the proliferative and apoptotic primary neurons from the E13.5 mice brain cortex. The primary neurons from the *Tyw1*−/− mice showed significantly decreased proliferation by using the EdU staining, while the intensity of apoptosis did not change significantly by using the TUNEL staining. *** p < 0.001, n.s.: no significant difference, unpaired t-test. Bar = 100 μm. **(G)** Quantification of the migrating primary neurons from the E13.5 mice brain cortex. In the transwell assay, the primary neurons from the *Tyw1*−/− mice showed significantly decreased migration. Red arrows indicated cells that already migrated through pores, while arrowheads indicated cells that were migrating in pores. *** p < 0.001, n.s.: no significant difference, unpaired t-test. Bar = 250 μm. **(H)** Schematics of the mechanism of ribosomal frameshift under the conditions of wild-type and mutant alleles of *TYW1*. GAA is the anticodon of tRNA^Phe^ for phenylalanine, and UUU is the codon of mRNA for phenylalanine. In the four mRNAs, with normalized lengths, of *SASS6*, *NCAPD2*, *CENPE*, and *STIL*, the positions of codon UUU were indicated by red bars. The introduction of mutated TYW1 led to promoted ribosomal frameshift at the interaction between tRNA^Phe^ and mRNAs, resulting in the reduced production of a subset of proteins involved in cell cycling. **(I)** The measurement of mRNA and protein expression of the E13.5 mice brain cortex. In the *Tyw1*−/− group, the RNA levels of *Cenpe*, *Stil*, *Ncapd2*, and *Sass6* did not change much compared to the WT group. However, the protein levels of Cenpe, Stil, Ncapd2, and Sass6 showed significant reduction in reference to the WT group, with remaining levels of 0.38, 0.63, 0.36, and 0.14, respectively. β-actin was used as the control in both RNA and protein measurement. Neither RNA nor protein of Tyw1 could be detected in the *Tyw1*−/− group. MW (Tyw1) = 84 kDa. MW (Cenpe) =312 kDa. MW (Stil) =143 kDa. MW (Ncapd2) =157 kDa. MW (Sass6) =74 kDa and 65 kDa. MW (β -actin) = 43 kDa. The quantitative RT-PCR and western blot data were from three independent experiments. * p < 0.05, *** p < 0.001, unpaired t-test.

Primary neurons extracted from the E13.5 mice brain cortex showed significantly decreased proliferation in the *Tyw1*−/− group compared to the wild-type group, while the apoptosis intensity showed no obvious increase in the *Tyw1*−/− group (Figure 4 (F)). The migration tests on the primary neurons proved reduced migrating capacity of the *Tyw1*−/− group compared to the wild-type one (Figure 4 (G)). In parallel, we evaluated the performance of SH-SY5Y cells after introducing *TYW1*-knockout (Supplementary figure 13). Interestingly, as observed in the primary mice neurons, the *TYW1*-knockout SH-SY5Y cells presented significantly reduced abilities of proliferation, adhesion, and migration, compared to the wild-type ones, however, there was no observable change in neuron differentiation (Supplementary figure 14).

To explore the mechanism of epigenetic regulation by *TYW1*, we constructed a model to simulate how the levels of wybutosine affected the ribosomal frameshift (Online methods). We defined a parameter (attenuation coefficient) for each protein to depict the vulnerability of a specific protein-coding gene to the reduced level of wybutosine. Typically, if the mRNA sequence of a specific gene does not contain a codon UUU, this gene will not be affected by the ribosomal frameshift related to the wybutosine level. On the contrary, if the mRNA sequence of a gene holds many codon UUU, its protein translation will be sensitive to the wybutosine level, with the extent determined by the density and distribution of codon UUU. This algorithm thus generated a protein list with attenuation coefficients lower than expectation, due to malfunctioning TYW1. Subsequently, we filtered the list by two thumb rules (Supplementary table 6). First, since we observed microcephaly and neurological manifestation in the subjects of human, mouse, and zebrafish, we chose proteins with low attenuation coefficients in all the three species. Second, we paid attention to proteins that were known to be disease-related, and bore resemblance to TYW1 in terms of gene expression patterns, especially in developing brains (Supplementary figure 15). In this way, four candidate proteins emerged, namely, CENPE (OMIM: 117143), STIL (OMIM: 181590), NCAPD2 (OMIM: 615638), and SASS6 (OMIM: 609321) (Figure 4 (H)). The measurement of E13.5 mice brain samples showed significantly reduced levels of these four proteins in the *Tyw1*−/− group compared to the wild-type one, while the corresponding RNA levels did not change much (Figure 4 (I)). Therefore, it was reasonable that the defective Tyw1 hindered the production of these four proteins in brain, with ensuing reduction of neuronal proliferation and migration. Interestingly, these four proteins are all involved in cell cycling and mitosis, the defects of which were reported to be related to primary microcephaly. STIL and SASS6 are necessary for centriole duplication during the cell cycle ^59–64^. CENPE is required for spindle microtubule capture and attachment at the kinetochore during cell division ^65–67^. NCAPD2 is a component of the condensin multiprotein complex participating in mitotic chromosome condensation ^68,69^.

### Null and hypomorphic alleles of *GPAM* incurred drastically reduced motor ability with unaffected cognition

The female proband CP_063_1, a spastic quadriplegic CP patient, was the only child of unrelated healthy parents without relevant family history. She was born by the Caesarean section at 39 gestational week when her mother was 30 years old and father was 32 years old. Her birth weight was 3.05 kg (−0.44 S.D.). The clinical examination at age of 2 years and 4 months showed body height 83 cm (−2.45 S.D.), weight 10 kg (−1.7 S.D.), and head circumference 47 cm (−1.31 S.D.). The cranial neuroimaging taken at 4 years old revealed no apparent anatomical abnormality except for plagiocephaly, which was due to the elongated period of supine position after birth. However, there was a reduction of white matter fiber tracts, especially in the corticospinal tract. The level of Gross Motor Function Classification System (GMFCS) was IV, meaning ‘self-mobility with limitations and may use powered mobility’. The fine motor skill was poor because the Manual Ability Classification System (MACS) was evaluated at level of III (handles objects with difficulty, need help to prepare and/or modify activities). Very interestingly, the cognitive evaluation (Wechsler Preschool and Primary Scale of Intelligence) was performed at age 3 years and 7 months, with full-scale IQ 116. The IQ scores ranked this CP patient above 70% of individuals in general population (IQ scores in a large unselected population follow a normal distribution with mean value 100 and S.D. 15). This patient epitomized a major subgroup, comprising half of the CP population, who were impaired in motion albeit normal in intelligence. The medical information of this patient was presented in Figure 5 (A) and Supplementary table 1 and 5.

**Figure 5:**
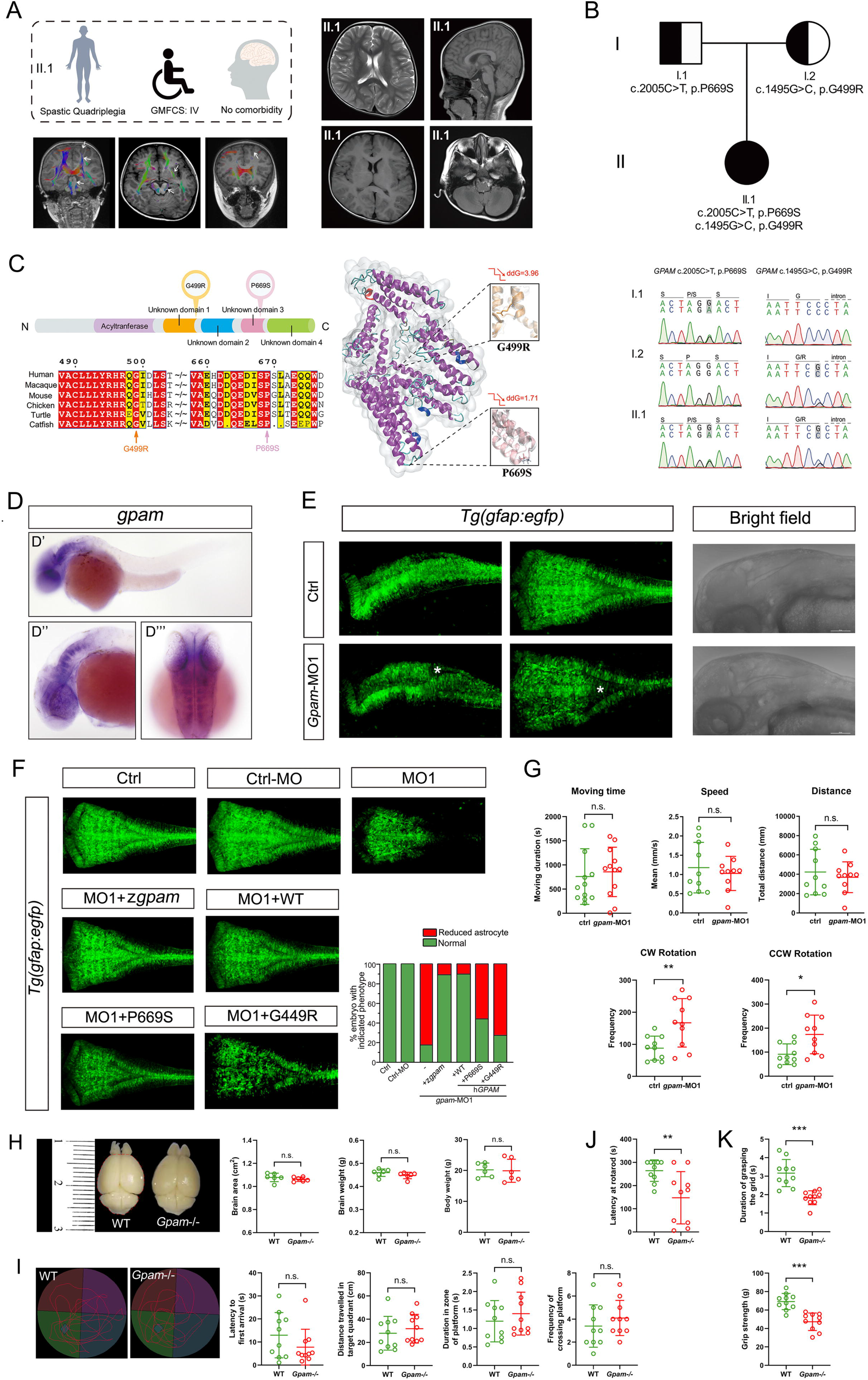
Correlation between phenotype and genotype of *GPAM*. **(A)** Brief medical description of the index patient II.1 (CP_063_1) harboring *GPAM* variants, and the results of magnetic resonance imaging (MRI) and diffusion tensor imaging (DTI). No gross anatomical abnormality was observed except for plagiocephaly. Deterioration of corticospinal tract was indicated by white arrows. **(B)** Compound heterozygous variants of *GPAM* were identified in the index patient, which co-segregated with the phenotype in the family members. **(C)** The two identified amino-acid changes, i.e., G499R and P669S, were both located inside the conserved protein domains. The two changes took over the amino-acid positions that were highly conserved among different species. Three-dimensional structural model of GPAM protein displayed the reduced protein stability (ddG, Gibbs free energy), and the local structural alterations before and after the introduction of two amino-acid changes. In the two zoom-in views, white bars stood for the wild-type amino acids and color bars for the mutated ones. **(D - G)** Morphological and behavioral assays on zebrafish models. **(D)** Whole mount *in situ* hybridization of zebrafish embryos at 36 hpf by using zebrafish antisense *gpam* probe showed specific expression of *gpam* in the brain, especially in the hindbrain. D’: lateral view. D”: lateral view with magnification of head region. D”’: dorsal view. hpf: hours post fertilization. **(E)** Confocal imaging of transgenic zebrafish *Tg(gfap:egfp)* embryos at 48 hpf showed reduced number of gfap^+^ cells in the *gpam* knockdown group (*gpam*-MO1, a splicing blocking morpholino), particularly in the hindbrain, labeled by white stars. Bright field lateral imaging showed no apparent general change in the brain. Left: lateral view. Middle: dorsal view. Right: lateral view. **(F)** The *gpam* knockdown (*gpam*-MO1) caused reduction of astrocyte in the zebrafish *Tg(gfap:egfp)* embryos at 48hpf, and there were only 17% of subjects that remained normal. This abnormal phenotype could be rescued by zebrafish *gpam* mRNA (*zgpam*), human *GPAM* mRNA (WT), mutated human *GPAM* leading to P669S, and mutated human *GPAM* leading to G449R, to variable extents (86%, 87%, 44%, 28%, respectively). Fish number ranged from 36 to 72 in different test groups. **(G)** Swimming behavior tests of the *gpam*-MO1 zebrafish larvae at 5 dpf compared to the controls, on moving time, speed, distance, clockwise (CW) rotation, and counterclockwise (CCW) rotation. The former three parameters showed no significant change, while the latter two parameters revealed significant changes. N = 10 to 12 for each test group. * p < 0.05, ** p < 0.01, n.s.: no significant difference, unpaired t-test. dpf: days post fertilization. **(H - K)** Morphological and behavioral assays on *Gpam* knockout (*Gpam*−/−) mice models. **(H)** The *Gpam*−/− mice at postnatal 8 weeks showed no significant alteration of brain size, brain weight, and body weight compared to the wild-type (WT) ones. The red dash line indicated the area of brain. N = 6 per genotype with equal number of male and female. n.s.: no significant difference, unpaired t-test. **(I)** Track plots depicted the trace of mice during the probe test in a Morris water maze. The *Gpam*−/− mice showed no difference in performance compared to the WT ones. N = 10 per genotype with equal numbers of male and female mice. n.s.: no significant difference, unpaired t-test. **(J)** Rotarod tests showed significantly reduced motor coordination and balance of the *Gpam*−/− mice compared to the WT ones. N = 10 per genotype with equal numbers of male and female mice. ** p < 0.01, unpaired t-test. **(K)** Grip strength tests revealed significantly shorter grip duration and weaker grip strength of the *Gpam*−/− mice compared with the WT mice. N = 10 per genotype with equal numbers of male and female mice. *** p < 0.001, unpaired t-test.

Compound heterozygous variants of *GPAM* (OMIM: 602395) were identified in this patient, the association between which and the CP phenotype in this family was confirmed by using targeted-PCR and Sanger-sequencing (Table 1 and Figure 5 (B)). The pRecessive value 0.9998 of *GPAM* (http://exac.broadinstitute.org/) rendered a very strong indicator for a recessive-disease-related gene. Allele frequencies of the two variants (NM_001244949: c.1495G>C, p.G499R; c.2005C>T, p.P669S) were both below 1 × 10^−5^, in the large-scale databases (https://db.cngb.org/cmdb/ and http://exac.broadinstitute.org/), and with no match in the in-house database of more than 2,247 healthy individuals. Pathogenicity prediction labeled the two variants damaging or disease-causing (Supplementary table 3). Multiple protein sequences alignment in a variety of species showed high conservation of both positions, i.e., G499 and P669, which were located inside the important protein domains. Three-dimensional structural models revealed that G499 and P669 were positioned at the pivotal regions, and both alterations of amino acids led to dramatic reduction of protein structural stability (Figure 5 (C)).

Using the whole-mount *in situ* hybridization (WISH) analysis of zebrafish, we found that *gpam* was specifically expressed in the developing central nervous system, mainly in the hindbrain (Figure 5 (D)). The morpholino-mediated *gpam* knockdown (*gpam*-MO) zebrafish models were established, by blocking mRNA splicing (MO1) and protein translation (MO2) of *gpam* (Supplementary figure 8 and 16). There was no apparent head size difference between *gpam*-MO zebrafish and the wild-type ones, but we observed obvious reduction of astrocyte in *gpam*-MO zebrafish, especially in the hindbrain (Figure 5 (E) and Supplementary figure 17). This reduction could be effectively rescued by the wild-type human *GPAM* or zebrafish *gpam* mRNA, but not by the *GPAM* mRNAs with variants causing G499R and P669S (Figure 5 (F)). This result corroborated the pathogenicity of the two variants identified in the patient. On the other hand, swimming behavior tests showed that the swimming capacity of *gpam*-MO zebrafish was compromised significantly (Figure 5 (G)), mimicking the motor impairment of the respective CP patient. Very interestingly, we noticed that the swimming capacity patterns of *tyw1*-MO and *gpam*-MO zebrafish were somehow complementary to each other, i.e., the swimming distance, speed, and moving time were abnormal in the *tyw1*-MO zebrafish but normal in the *gpam*-MO zebrafish, in contrast, the rotation movement was normal in the *tyw1*-MO zebrafish but abnormal in the *gpam*-MO zebrafish. This finding may reflect the different underlying mechanism posed by the two genes.

Meanwhile, we constructed *Gpam* knockout mice models by using CRISPR/Cas9 genome editing technology (Supplementary figure 11). In *Gpam* knockout (*Gpam*−/−) mice, there was no significant alteration of brain size or weight compared to the wild-type littermates (Figure 5 (H)), compatible with the observation in human and zebrafish. Similarly, the Morris water maze tests showed no significant difference between the *Gpam*−/− mice and the wild-type ones (Figure 5 (I)). However, the rotarod test and the grip strength test revealed significant reduction of the motor capacity in the *Gpam*−/− mice compared to the wild-type ones (Figure 5 (J) and (K)).

### Defective Gpam interfered with proliferation of astrocyte and oligodendrocyte myelination by perturbing lipid metabolism

The gene of *GPAM* encodes a mitochondrial enzyme that catalyzes the production of lysophosphatidic acid (LPA) by using glycerol-3-phosphate (G3P) as the substrate, which is the rate-limiting step in the synthesis of glycerolphospholipids and triacylglycerol (TAG) ^70^. There are four isoenzymes in this step, among which GPAM occupies most of the activity in the liver, and the hepatic knockdown of Gpam in mice reduced the levels of TAG, diacylglycerol, fatty acid and cholesterol ^70–72^. In the central nervous system, GPAM was specifically expressed in astrocyte, and reached highest expression level on P14 in mice hindbrain (Figure 2 (D) and (E); Figure 6 (A) and (B)). The enzymatic activity in astrocyte is maintained mainly by GPAM, supplemented by GPAT4, an isoenzyme ubiquitously expressed in a variety of brain cells including neurons and oligodendrocyte ^73,74^.

**Figure 6:**
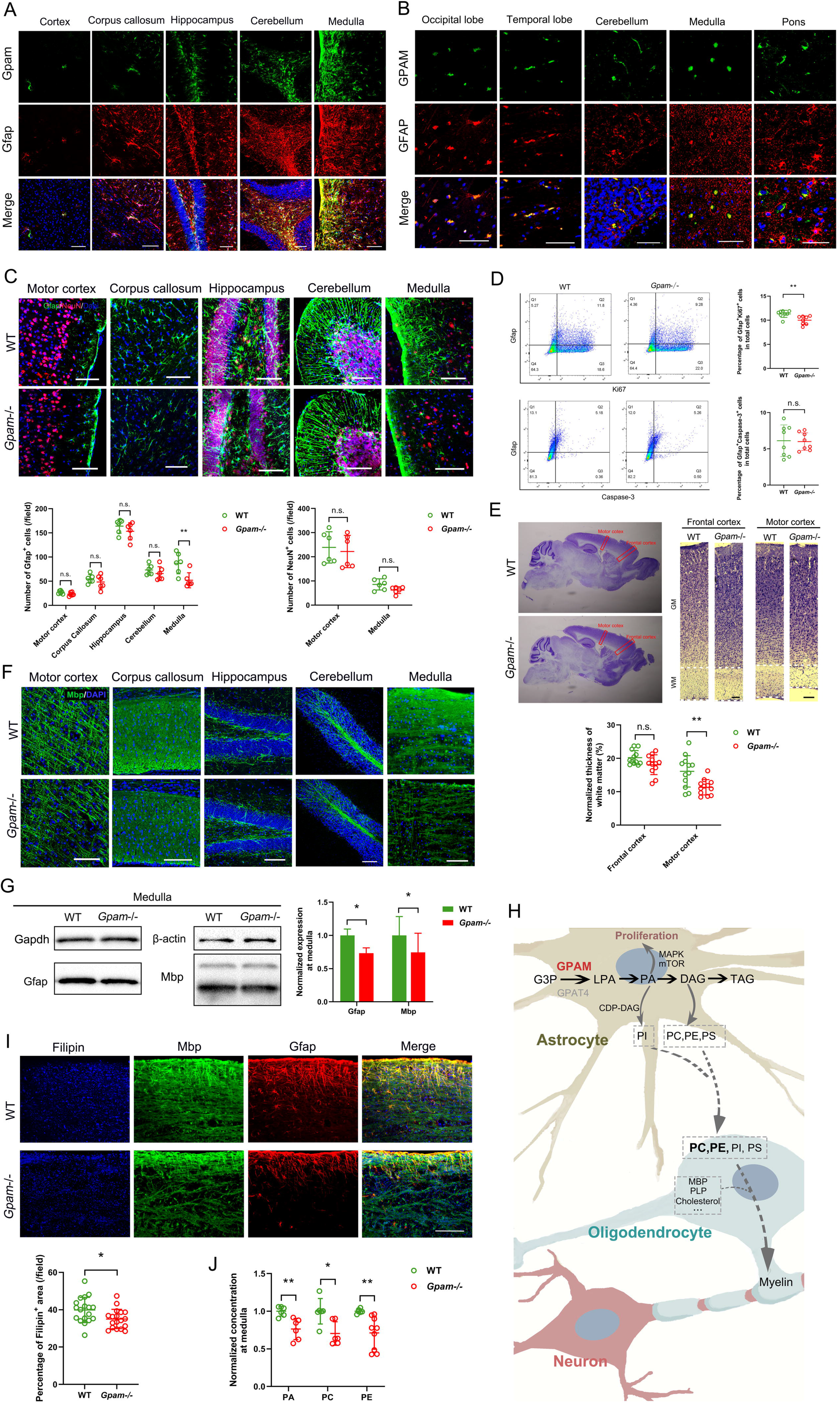
Functional mechanism of defective GPAM. **(A)** Brain section immunostaining of the wild-type mice at postnatal 8 weeks, where Gpam (green) and Gfap (red) were co-labeled, showed the Gpam expression in the astrocyte of many brain regions. Blue: DAPI. Bar = 100 μm. **(B)** Immunostaining of the normal human brain tissue microarrays, where GPAM (green) and GFAP (red) were co-labeled, revealed the GPAM expression in the astrocyte of a variety of brain regions. Blue: DAPI. Bar = 50 μm. **(C)** Brain sections immunostaining of the *Gpam*−/− mice at postnatal 8 weeks revealed significantly reduced number of the Gfap+ cells in medulla, compared to the wild-type (WT) ones. The same tendency was observed in other brain regions albeit without statistical significance. Green: Gfap. Red: NeuN. Blue: DAPI. Bar = 100 μm. N = 3 per genotype. For a specific sample, each measurement was performed twice in separate fields. ** p < 0.01, n.s.: no significant difference, unpaired t-test. **(D)** Flow cytometry of whole brain cells lysis of P1 mice stained with Gfap, Ki67, and Caspase-3. Percentages of the Gfap^+^Ki67^+^ cells were significantly lower in the *Gpam*−/− mice brain compared to the WT ones, indicating inhibited proliferation of astrocyte. Percentages of the Gfap^+^Caspase-3^+^ cells showed no difference between the two groups, indicating no activated apoptosis. N = 8 per genotype with equal numbers of male and female mice. ** p < 0.01, n.s.: no significant difference, unpaired t-test. **(E)** Nissl staining of brain sagittal sections of mice at postnatal 8 weeks showed a significantly thinner white matter at the motor cortex of the *Gpam*−/− group, compared to the WT ones. The same tendency was observed at the frontal cortex albeit without statistical significance. Bar = 200 μm. N = 3 per genotype. For a specific sample, each measurement was performed four times in separate fields. ** p < 0.01, n.s.: no significant difference, unpaired t-test. **(F)** Brain sections immunostaining of mice at postnatal 8 weeks showed lower density of the myelin binding protein (Mbp) at medulla of the *Gpam*−/− mice, compared with the WT ones. Green: Mbp. Blue: DAPI. Bar = 100 μm. **(G)** Western blot of mice brain medulla showed significant reduction of Gfap and Mbp in the *Gpam*−/− group compared to the WT one. Gapdh and β -actin were used as controls. MW (Gfap) = 50 kDa. MW (Mbp) = 19 and 26 kDa. For a specific sample, measurement was performed three times independently. * p < 0.05, unpaired t-test. **(H)** Schematics of the mechanism of GPAM involvement in lipid metabolism and its influence on astrocyte proliferation and oligodendrocyte myelination, by regulating the production of PA, PI, PC, PE, PS, etc. GPAT4: glycerol-3-phosphate acyltransferase 4. G3P: glycerol-3-phosphate. LPA: lysophosphatidic acid. PA: phosphatidic acid. DAG: diacylglycerol. TAG: triacylglycerol. CDP-DAG: cytidine diphosphate diacylglycerol. PI: phosphatidylinositol. PC: phosphatidylcholine. PE: phosphatidylethanolamine. PS: phosphatidylserine. MBP: myelin binding protein. PLP: proteolipid protein. **(I)** Brain medulla sections immunostaining showed significant reduction of Filipin, as well as Mbp and Gfap, in the *Gpam*−/− mice compared to the WT ones. Percentage of the Filipin^+^ area per imaging field was proportionate to the cholesterol abundance. Bar = 100 μm. N = 3 per genotype. For a specific sample, each measurement was performed four times in separate fields. * p < 0.05, unpaired t-test. **(J)** Quantification of PA, PC, and PE levels at the brain medulla of mice at postnatal 8 weeks revealed significant reduction in the *Gpam*−/− group compared to the WT one. N = 3 per genotype. For a specific sample, each measurement was performed twice or thrice. * p < 0.05, ** p < 0.01, unpaired t-test.

In the *Gpam*−/− mice, we observed the reduced number of astrocyte, especially in the medulla regions (Figure 6 (C)). The whole brain cell lysis of P1 mice showed significantly reduced proliferation of astrocyte in the *Gpam*−/− mice, but no apparent increase in the intensity of apoptosis was observed (Figure 6 (D)). Interestingly, brain section staining revealed an obviously decreased white matter thickness of the motor cortex in the *Gpam*−/− mice (Figure 6 (E)). In parallel with the decrease of astrocyte density, a myelin marker, namely, myelin binding protein (Mbp), was revealed to be significantly reduced in the medulla regions of the *Gpam*−/− mice brain (Figure 6 (F) and (G)). Given the manifestation and cranial MRI results of the respective CP patient, we assumed that, the phenotype of the *Gpam*−/− mice should be attributed to the hypomyelination of corticospinal tract that originates from the primary motor cortex, goes through and modulates in the medulla, and ends in the spinal cord ^75^.

It is known that myelin is a specialized form of oligodendrocyte membrane that tightly wraps and insulates the axons to accelerate the action potential conduction ^48,76^. Compared with other membranes, myelin has a much higher content of lipid, up to 80% of the total dry weight, and the most abundant lipid groups in myelin are cholesterol, phospholipids, and glycolipids ^76–78^. It was reported that in mice models, lipid synthesis blockage in the oligodendrocyte caused instant onset of severe demyelination and neurological symptoms, however, this damage could be spontaneously remedied to much extent after a period of months ^79^. Since the blood brain barrier shields the central nervous system from circulating lipids, an exogenous lipid supply in the parenchyma of brain should have accounted for this phenomenon, and fortuitously, recent studies discovered that the major supplier of lipids in brain was astrocyte ^80–82^. Actually, it was revealed that the horizontal lipid flux from astrocyte to oligodendrocyte was a major feature of myelination, and disruption of lipid synthesis in astrocyte led to lasting and more severe hypomyelination than inactivation of that in oligodendrocyte ^83–85^. In astrocyte, GPAM is committed to a rate-limiting step from G3P to LPA, the subsequent lipid products of which include phosphatidic acid (PA), diacylglycerol (DAG), triacylglycerol (TAG), phosphatidylinositol (PI), phosphatidylcholine (PC), phosphatidylethanolamine (PE), phosphatidylserine (PS), among others. PI, PE, PC, and PS are the major components of myelin ^86,87^ (Figure 6 (H)). Noticeably, a large body of evidence implicated the association between *GPAM* polymorphisms and plasma level of cholesterol ^88–93^. Experiments on brain sections and extraction confirmed this hypothesis, with significantly reduced contents of lipids detected at medulla of the *Gpam*−/− mice (Figure 6 (I) and (J)). Interestingly, we also observed a significant reduction of 1,2-diacyl-G3P (PA) from the samples, which is a downstream product of GPAM-involved reaction (Figure 6 (H) and (J)). PA is not only a precursor of phosphatidylinositol (PI), a component of myelin, but also promotes cell proliferation in astrocyte by activating the MAPK and mTOR signaling pathways (Figure 6 (H)) ^94–96^. Inhibition of PA was known to reduce the number of astrocyte and cause neurodevelopmental disorder^97–100^.

## Discussion

This work is, to the best of our knowledge, the first large-scale, full-dimensional, clinical genetics research on the etiology of cerebral palsy. We identified, in 45% of 120 CP families, detrimental variants of the established and novel genes related to CP and other NDDs. The meta-analysis on the 117 CP-related genes found in this study and garnered from literature, created an unprecedented compendium spanning a variety of attributes of CP-related genes. Intriguingly, we came up with a novel classification system for CP-related genes, according to their dichotomous spatiotemporal expression patterns. This can serve as a useful guideline for exploring the underlying mechanism and designing targeted treatment.

In parallel with this research, we managed to take full use of the etiological information in the clinical practice. The families were informed of the prognostic outcomes based on gene-wise knowledge bases, and received the adjusted medical and rehabilitation treatments, plus genetic counseling as they wished. For example, we offered serine and glycine replacement to CP_035_1 with *PSAT1* defects ^101^, Doconexent to CP_044_1 with *CYP2U1* defect ^102,103^, and magnesium carbonate to CP_053_1 with *CNNM2* defect ^104^. Likewise, we implemented personalized rehabilitation training for countering the specific progression of motion disability (Figure 1 and Supplementary table 1).

*TYW1* is a novel CP-related gene, defects of which were confirmed in this study to result in primary microcephaly and deterioration of both motion and cognition, by disturbing proliferation and migration of neurons in the developing brain, including motor cortex and frontal cortex. Blockage of neuronal proliferation and/or migration during brain development is a common theme in etiology of primary microcephaly ^105,106^. Interestingly, most of the known MCPH, i.e., microcephaly primary hereditary, genes encode components of machinery of cell division and cell cycling, while reports on genes like *TYW1* that is in charge of a subset of MCPH genes, are rare, if not absent. TYW1 plays a critical role in generating wybutosine, a hypermodification at position 37 of tRNA^Phe^ responsible for stabilizing codon-anticodon binding, and this epigenetic procedure can be grouped into one of the three aspects of tRNA activation, namely, the biogenesis, charging, and modification of tRNAs ^56,57^. We noticed that, a large proportion of genes involved in tRNA charging have been found to be associated with disease, especially NDDs, however, merely ~10% of genes participating in tRNA modification are related to human illness ^54^. Because of the enduring existence of most tRNA modification throughout species evolution, we believed that tRNA modification may be underestimated in its role in disease onset and progression. Future studies in this field can be modeled after this work on *TYW1*, and since the great majority of tRNA modification were identified at position 34 and 37 in the anticodon stem loop, the algorithm that we developed for wybutosin at postion 37 may find a multitude of scenarios to be applied to.

*GPAM* encodes a mitochondrial enzyme specifically expressed in astrocyte and plays a crucial role in lipid metabolism in brain. Aberrant GPAM found in this study reduced a variety of lipid contents in astrocyte and oligodendrocyte, causing hypomyelination in corticospinal tract and suppressing proliferation of astrocyte *per se*, and the latter deteriorated the interplay between oligodendrocyte and astrocyte thus aggravated the hypomyelination ^85,107^. Lipid metabolism problems resulted in cases of neurodevelopmental disorders, including ID, ASD, attention deficit hyperactivity disorder, and schizophrenia ^108–112^. And genetic defects in astrocyte were known to be related to lesions in brain white matter, such as *GFAP* and *NPC1* ^113–116^. This work disentangled a complex mechanism between astrocyte and myelination, providing a chance to develop new therapeutic strategies. Very intriguingly, there were reports in mice models that, high-fat diet could promote circulating lipid contents and ameliorate hypomyelination in brain if the lipid synthesis in astrocyte was blocked; therefore, it is tantalizing to apply this to the *Gpam*−/− mice and human patients with similar lipid metabolic problems ^84,117^.

## Supporting information

Supplementary table 1 CP cohort medical chart

Supplementary table 2 Deep sequencing performance

Supplementary table 3 Details of the CP-related variants and incidental findings

Supplementary table 4 Known CP-related genes

Supplementary table 5 Medical charts of the patients with TYW1 and GPAM defects

Supplementary table 6 Predicted attenuation coefficients of proteins under the conditions of TYW1 knockout

## Data availability

This study’s samples and data are accessible, conforming to the principles of the International Cerebral Palsy Genomics Consortium (ICPGC) (https://icpgc.org/).

## Acknowledgement

We are very grateful to the families participating in this study. This work was supported by the Major Medical Collaboration and Innovation Program of Guangzhou Science Technology and Innovation Commission (201604020020), the National Natural Science Foundation of China (81671067, 81974163, 81672253, 31871063, and 81701451), the China Postdoctoral Science Foundation (2019M662852), the Natural Science Foundation of Guangdong Province of China (2018A030313538), and the Key-Area Research and Development Program of Guangdong Province (2019B020227001).

## Author contribution

H.H. conceived and coordinated the project. K.X., H.T., and L.H. recruited the cohort. D.L., J.Z., and X.W. performed the zebrafish modeling. D.H. and N.L. oversaw the cell and mice modeling with aids of P.Z., H.T., Y.Q., C.S., F.Q., M.Y., Z.Z., C.L., S.Z., Y.L., L.L., J.H., and C.P‥ N.L. was in charge of the genetic analysis and bioinformatics with aids of D.H., X.F., Y.L., X.P., Z.Z., T.X., Q.Z., and L.L‥ N.L. and H.H. wrote the manuscript with comments from all authors.

## Online methods

### Cohort recruitment and sample preparation

#### Cohort recruitment

The family history and clinical characteristics of CP individuals were documented regarding the CP phenotypic profiles. The motor signs (spastic, ataxic, dyskinetic, etc.) and anatomical distribution (monoplegia, diplegia, triplegia, hemiplegia, and quadriplegia) of movement disorder were evaluated and categorized. Motor severity was measured by the Gross Motor Function Classification System (GMFCS) on 5 levels ^1^. The common co-morbidities, i.e., ID/ASD, vision/hearing impairment, epilepsy, dyslexia, etc., were recorded for each patient. The fine motor skill was evaluated by the Manual Ability Classification System (MACS) ^2^. Rehabilitation training outcome was determined by the Activity of Daily Living (ADL) ^3^, and the Gross Motor Function Measure (GMFM) ^4^, both of which were measured twice before and after treatment.

#### Magnetic Resonance Imaging (MRI) and diffusion tensor imaging (DTI)

All MRI imagings were performed on a 3.0T Siemens scanner (Magnetom Systems, Skyra, Munich, Germany). For DTI, an echoplanar sequence with diffusion gradients (b = 1000 s / mm^2^) applied in 64 noncollinear directions was used. Basic parameters included: TR (10000 ms) / TE (92 ms), Voxel (2.0 × 2.0 × 2.0 mm), FOV (256 × 256), layer thickness (2.0 mm). DTI measurement was three-dimensional reconstruction of color-coded FA graphics through DTI Studio (DWI standard deviation).

#### Sample preparation

Genomic DNA from all the members in the CP cohort was extracted from the peripheral blood by using the QIAamp DNA blood mini kit (Qiagen, Frankfurt, Germany) according to the manufacturer’s instructions.

PCR amplification was performed by using the Q5 High-Fidelity Polymerase (5X, NEB, MA, USA) with the specific primers targeted to the selected candidate variants listed as below:

*TYW1*-R206C-Forward: 5’-TTTCCTCTTCCTTAGCGGCA-3’,
*TYW1*-R206C-Reverse: 5’-CAATCCTCCCAACTCCACCT-3’;
*TYW1*-R389Q-Forward: 5’-CTCCAGGATGCATTCCCTTA-3’,
*TYW1*-R389Q-Reverse: 5’-TCGATCACCATGACCTTTCA-3’;
*GPAM*-P669S-Forward: 5’-GAGGGGTGCAGGTGTAGGTA-3’,
*GPAM*-P669S-Reverse: 5’-AGGAGATGGCCTGGAAAAGT-3’;
*GPAM*-G499R-Forward: 5’-CTGAGAACCCCAGGTCAAAA-3’,
*GPAM*-G499R-Reverse: 5’-TGATGGAAGAGACCACACCA-3’.

Then, the PCR products were processed by the Sanger Sequencing to determine the carrier status of the specific variants.

### Fragile-X screening and mitochondrial genome sequencing

#### Fragile-X screening

We performed the fragile X screening for the gene of *FMR1* on 120 families including patients, unaffected siblings, and parents. Fragile-X syndrome (FXS) is the most common monogenic form of intellectual disability and autism spectrum disorder, predominately due to the expansion of a CGG repeat located at the 5’ UTR of the *FMR1* gene (OMIM: 309550). Expansion to the full mutation (more than 200 repeats) leads to hypermethylation and silencing of *FMR1*, resulting in the absence of its protein product, FMRP, which causes FXS. A *FMR1* CGG repeat was amplified with the primers (Forward: 5’-GCGCTCAGCTCCGTTTCGGTTTCACTTCC-3’ and Reverse: 5’-CCCAAGTCCAGTCCTTCCCTCCCAACAACA-3’) by using the LA Taq polymerase kit (RR02BG, TaKaRa, Shiga, Japan) and 50 ng of total genome DNA, then the PCR products were analyzed by gel electrophoresis and Sanger sequencing. Thermal cycling was as follows: denaturation at 96 °C for 3 min and 10 cycles of 98 °C for 20 sec, 65 °C for 45 sec, and 72 °C for 3 min, followed by 22 cycles of 98 °C for 20 sec, 68 °C for 3.5 min, and a final extension at 68°C for 10 min ^5^. There is no full mutation observed in this cohort.

#### Mitochondrial genome sequencing

We conducted very-high-coverage (6000X) mitochondrial DNA sequencing for all CP patients. The 16.6 kb circular mitochondrial genome encodes 13 protein subunits of the electron transport chain, 22 tRNA and 2 rRNA genes ^6^. The mitochondrial genome was amplified by a single-amplicon LR-PCR using a LA Taq polymerase kit (RR02BG, TaKaRa, Shiga, Japan) and 100 ng of total genome DNA ^7^. Indexed paired-end DNA libraries were prepared and sequenced by Nextseq500 sequencer (Illumina, CA, USA), according to the manufacturer instructions. The sequenced data was analyzed by the NextGENe (SoftGenetics, State College, PA) with the mitochondrial reference sequence (NC_012920). The average sequencing coverage of mitochondrial genome was 6000X, and the variants with abundance >2% were reported, because of the well-known heteroplasmy of mitochondrial variants. The pathogenicity of a specific variant was defined according to the guideline recommended by the American College of Medical Genetics and Genomics (ACMG) ^8^. The pathogenic mitochondrial variants were validated by PCR and Sanger sequencing of the total DNAs from the patients and the biological mothers.

### High-density cytogenetic microarray

High-density cytogenetic microarrays for all CP patients were performed by using the CytoScan HD Array platform (Affymetrix, CA, USA) with more than 2.6 million markers including 0.75 million single nucleotide polymorphisms. CNV detection was done by using the Chromosome Analysis Suite (version 3.0, Affymetrix, CA, USA). The genomic coordinates were based on the Human Genome Build GRCh37/hg19. Pathogenic CNVs and likely pathogenic CNVs were defined as the rules recommended by the American College of Medical Genetics and Genomics guidelines (ACMG) ^9^. CNVs were compared in the Database of Genomic Variants (http://dgv.tcag.ca/dgv/app/), CAGdb (Cytogenomics Array Group Database) (http://www.cagdb.org/), ISCA (International Standards for Cytogenomic Arrays) (http://dbsearch.clinicalgenome.org/), DECIPHER (https://decipher.sanger.ac.uk/), and an in-house 2,247 control samples. Pathogenic and likely pathogenic CNVs were confirmed in the patients and parental samples using a SYBR Green-based real-time quantitative PCR assay (A25778, Thermo Fisher Scientific, MA, USA).

### Whole genome sequencing and ultra-high-depth whole exome sequencing

Whole genome sequencing (WGS) was conducted for all 385 members of the 120 CP families including 122 patients, their parents and unaffected siblings, on a HiSeq XTen Deep Sequencer (Illumina, CA, USA), with an average coverage of ~36X, according to the manufacturer’s manuals. The primary data analysis was done by a combination of BWA sequence aligner for reads alignment to human reference genome GRCh37/hg19, GATK pipeline for calling single-nucleotide variant (SNV) and small insertion and deletion (indel), and ANNOVAR for variant characterization ^10–12^.

The 66 patients with no obvious causal variant underwent further assessment with the ultra-high-depth whole-exome sequencing (UWES) in order to capture post-zygotic mutation (PZM) and those variants located in poorly-represented coding regions in WGS. The UWES procedures were performed on a HiSeq4000 deep sequencer (Illumina, CA, USA) with a 150bp paired-end protocol after enrichment in the coding regions by the SureSelect Human All Exon V5 kits (Agilent, CA, USA). The average coverage of UWES for CP patients reached more than 600X. The generated raw sequences were processed by the MERAP package for alignment, quality control, calling single nucleotide variant (SNV), insertion and deletion (Indel), structural variation (SV), and copy number variation (CNV), plus the variants annotation and prioritization ^13^.

The sequencing performances of WGS and UWES were recorded in Supplementary table 2.

### Analysis pipeline of deep sequencing data

The variant prioritization and evaluation of etiologic involvement were performed by two pipelines, namely, MERAP and ANNOVAR. In brief, MERAP pipeline first filters all identified variants through comparison with the disease-associated variants in the Human Gene Mutation Database (HGMD, 2020.2) and the Online Mendelian Inheritance in Man (OMIM) to collect those known disease-causing variants. In order to filter out neutral variants, MERAP uses entries from the dbSNP143 (http://www.ncbi.nlm.nih.gov/projects/SNP/), 1000 Genome (http://www.1000genomes.org/), NHLBI Exome Sequencing Project (ESP, http://evs.gs.washington.edu/EVS/), ExAc (http://exac.broadinstitute.org/), and the Chinese Millionome Database (CMDB) (https://db.cngb.org/cmdb/) as screening databases. In principle, candidate variants causing recessive traits should not occur in healthy controls as homozygotes, and the frequency of respective heterozygotes should not exceed 0.1%. MERAP uses the RefSeq genes (http://www.ncbi.nlm.nih.gov/refseq/) as reference, and nonsynonymous changes are described in terms of gene ID, base change, protein change, genomic coordinate, transcript coordinate, protein coordinate, protein length, affiliated with gene description from the Human Gene Nomenclature Committee (HGNC, http://www.genenames.org/). Changes destroying conventional splice sites or introducing novel splice sites are identified by MERAP’s module called SSFinder. To assess the pathogenicity of missense mutations, MERAP generates a single score integrating the results of seven different algorithms, including the Grantham score, PhyloP, GERP, SIFT, PolyPhen2, Mutation-Taster, and the Conserved Domains Database (CDD, http://www.ncbi.nlm.nih.gov/Structure/cdd/cdd.shtml). With empirical false discovery rate cutoffs, this score serves as dichotomized pathogenicity predictions even if any two of the seven algorithms might not coincide, as is often the case. MERAP rules out candidate genes reported to harbor homozygous loss-of-function (LOF) variants in healthy individuals, which applies to >1% of the human genes. Typically, if more than three independent truncating variants are observed in >10 of the exomes listed in the 1000 Genome, Exome Variant Server (ESP6500), NHLBI GO Exome Sequencing Project (ESP) (http://evs.gs.washington.edu/EVS/), and ExAC (http://exac.broadinstitute.org/) databases, the relevant gene is flagged as LOF tolerant. To facilitate the choice between few remaining candidate genes, MERAP also provides a list of ~4,500 known disease genes extracted from OMIM (http://www.ncbi.nlm.nih.gov/omim), ClinVar (http://www.ncbi.nlm.nih.gov/clinvar/), and HGMD (http://www.hgmd.org/), as well as more than 8,000 associated disorders and their symptoms. For novel candidate genes without prior link to disease, MERAP offers information on their interaction with known disease genes, based on data from Biogrid (http://thebiogrid.org/) and IntAct (http://www.ebi.ac.uk/intact/), assuming that genes implicated in clinically similar disorders tend to cluster in gene or protein interaction networks. Variants were considered *de novo* if neither parent had the variant, and candidate variants were selected by segregation analysis. The pathogenic and likely pathogenic genes/variants were defined according to the standards and guidelines of ACMG ^8^.

The ANNOVAR pipeline was also used, with additional considerations from the Residual Variation Intolerance Score (RVIS) and the Combined Annotation-Dependent Depletion (CADD) score ^11^. The former ranks genes in terms of intolerance to functional genetic variation and the latter integrates several well-known tools. We empirically set a cutoff RVIS score of 50th percentile for known and novel genes and a cutoff CADD score of 20 for novel candidate genes. When defining likely causal variants/genes, we followed the guidelines designated by ACMG.

All variants of putative clinical relevance were confirmed by the conventional PCR and Sanger sequencing. Parent-child relationships were confirmed by using the PLINK with SNPs drawn from WGS which matched the CytoScan SNP repertoire ^14^. The false positive rate of WGS was generated by the comparison between Sanger sequencing and WGS, and we found no false positive variant called by our pipeline. The false negative rate of WGS was evaluated by the SNP genotyping comparison between WGS and CytoScan high-density microarray, which resulted in an average false negative rate of less than 0.1%.

### Gene-set enrichment, pathway analysis, gene expression analysis and protein-protein interaction analysis

#### Gene ontology and KEGG pathway analysis

Gene ontology analysis was performed on 117 CP-related genes and 706 NDD-related genes ^15^ separately using GO enrichment analysis tool of Omicshare (https://www.omicshare.com/tools/home/report/goenrich.html). The top 27 GO terms with p-value < 0.01 were selected and presented. To NDD-related genes, the same GO terms were filtered with p-value < 0.01, and only those with rich factor > 0.09 (for Biological process and cellular component) and 0.07 (for Molecular function) were indicated.

KEGG pathway analysis was performed on 117 CP-related genes and 706 NDD-related genes ^15^ separately using pathway enrichment analysis tool of Omicshare (https://www.omicshare.com/tools/home/report/koenrich.html). The top 27 pathways with p-value < 0.05 were selected and presented. To NDD-related genes, same pathways were filtered with p-value < 0.05, and only those with rich factor (the ratio between number of genes from the present gene-set and all genes enriched in a certain pathway) > 0.1 were indicated.

#### Expression analysis of CP genes

To visualize the expression pattern of CP-related genes in different brain regions, we extracted the bulk RNA-seq data of CP-related genes from the published data of Li *et al.* ^16^. On the other hand, to investigate the expression profile of CP-related genes in different cell types, we used the R package Seurat to import, filter, normalize and scale (with default settings) the single cell RNA-seq data for human brain obtained from the published data of Li *et al.* ^16^ and Lake *et al.* ^17^. The expression level of CP-related genes in different brain regions and different cell types were displayed in the form of heatmaps by using TBtools software ^18^.

#### Protein-protein interaction network

Protein-protein interaction network between CP-related genes and NDD-related genes was obtained from the STRING database ^19^ (https://string-db.org/). The interaction network model was generated with Cytoscape ^20^.

### Homology modeling

Domains of TYW1, GPAM, CDK5RAP1, PTK7 and TARS were predicted by the InterProScan protein domains identifier ^21^ (http://www.ebi.ac.uk/interpro/search/sequence-search). For GPAM, PTK7 and TARS, unknown domains were also predicted with the MEME (Multiple EM for Motif Elicitation) motif discovery tool ^22^ (http://meme-suite.org/tools/meme). Multiple sequence alignments of TYW1, GPAM, CDK5RAP1, PTK7 and TARS were generated by using the ClustalW program ^23^.

Three-dimensional structural models of TYW1 (TYW1_WT, TYW1_R206C and TYW1_R389Q), GPAM (GPAM_WT, GPAM_G499R and GPAM_P669S), CDK5RAP1 (CDK5RAP1_WT, CDK5RAP1_M255T and CDK5RAP1_E494V), PTK7 (PTK7_WT, PTK7_G476R and PTK7_P698R) and TARS (TARS_WT, TARS_T258S and TARS_Q702R) were predicted by the I-TASSER web tool ^24^ (http://zhang.bioinformatics.ku.edu/I-TASSER/). Visual representation of models and structural superposition were generated by a software package named visual molecular dynamics (VMD) ^25^.

The mutant stability change (∆∆G) of variants of TYW1 (TYW1_R206C and TYW1_R389Q), GPAM (GPAM_G499R and GPAM_P669S), CDK5RAP1 (CDK5RAP1_M255T and CDK5RAP1_E494V), PTK7 (PTK7_G476R and PTK7_P698R) and TARS (TARS_T258S and TARS_Q702R) were predicted by using the STRUM server ^26^ (https://zhanglab.ccmb.med.umich.edu/STRUM/).

### SH-SY5Y cell modeling

The human neuroblastoma cell line SH-SY5Y was retrieved from the China National Infrastructure of Cell Line Resource (resource code: 3131C0001000700097).

#### CRISPR/Cas9-mediated *TYW1* gene knockout in SH-SY5Y cells

Gene knockout was performed in SH-SY5Y. Briefly, all-in-one CRISPR/Cas9 system was used to produce lentivirus expressing the single guide RNA (sgRNA) and Cas9 protein (U6-sgRNA-EF1α-Cas9-P2A-Puro). The sgRNA sequence is 5’-GCATCGTGTGATGAGTCGAG-3’ for *TYW1* (Edigene, Beijing, China). Sequences were selected with online sgRNA design tool CRISPOR (http://crispor.tefor.net/). SH-SY5Y cells were lentiviral transduced. After 48 h, cells were selected with puromycin at a concentration of 3.5 mg/ml.

Genome editing status in the mixed clonal KO cells was determined by Sanger sequencing analysis with the TIDE web tool ^27^. Knockout efficiency was confirmed by RT-qPCR using primers as followed:

*TYW1*-locus1-Forward: 5’-TGCAGGTGGACAAAGTCC-3’,
*TYW1*-locus1-Reverse: 5’-TTATTAGCACACGCCAAGC;
*TYW1*-locus2-Forward: 5’- CAGACTGGAACAGCGAAGGGA-3’,
*TYW1*-locus2-Reverse: 5’-TGTATGTCGCAACCAGGAAGA-3’;
*ACTB*-Forward: 5’-TGCGTGACATTAAGGAGAAG-3’,
*ACTB*-Reverse: 5’-CATGATGGAGTTGAAGGTAGT-3’.

#### Analysis of cell viability and adhesion

Cell viability was examined with CCK-8 detection kit (Doindo, kumamoto, Japan). Cells in each group were seeded into 96-well plate at a density of 5 × 10^4^ cells/well. After 12 h, medium was removed. 100 μl CCK-8 working solution (1:10 in medium) was added in each well. Cells were incubated for 4 h. Cell viability which reflected cell proliferation was detected by OD_450_ with a microplate reader.

In adhesion experiment, 96-well plates were coated with matrigel (Corning, NY, USA). SH-SY5Y cells in the wild-type and *TYW1*-KO groups were both divided into control part and experiment part. Medium was added in background part. Plates were incubated at 37 °C for 1 h. In experimental part, cells were washed three times by using washing buffer. Then, staining solution (1:10 in medium) was added in control part, experimental part and also in background part. After 4 h, OD_450_ was measured. The adhesion rate was calculated as following: the adhesion rate of cells (%) = (OD_450_ of experimental part – background) / (OD_450_ of control part-background) × 100. Measurement was recorded on three independent experiments.

#### Detection of cell migration

Cells in each group were seeded on 6-well plates at a density of 2 × 10^6^ cells/well. After reaching 90% confluence, a straight scratch wound was made using a pipette tip. Then, cells were washed with PBS to remove suspended cells gently and added with new medium. At 0 h, 24 h and 48 h, the wound area was observed under microscope. At 48 h, wound in the wild-type group almost closed. The cell migration was quantified according to the wound area at 48 h relative to the initial wound area by the ImageJ software. Measurement was recorded on three independent experiments.

#### Induction of cell differentiation

Cells in each group were seeded on 24-well plates coated with matrigel (Corning, NY, USA). After 24 h, 10 μM all-trans retinoid-acid (RA) (Stemcell technologies, Vancouver, Canada) was added to induce the cell differentiation. At day 5 in the presence of RA, cells were treated with 50 ng/ml BDNF (Stemcell technologies, Vancouver, Canada) in DMEM without serum. Morphology of cells was observed by using immuno-staining of MAP2 at day 10. Neurites were traced with the NeuroGrowth plugin of ImageJ software to examine the number and total length of branches. Measurement was recorded on three independent experiments.

### Zebrafish modeling

All zebrafish experimentation was carried out in accordance with the NIH Guidelines for the care and use of laboratory animals (http://oacu.od.nih.gov/regs/index.htm) and ethically approved by the Administration Committee of Experimental Animals, Jiangsu Province, China [Approval ID: SYXK (SU) 2007–0021].

#### Zebrafish lines and whole mount *in situ* hybridization

The zebrafish embryos and adults were maintained in the Zebrafish Center of Nantong University under conditions in accordance with our previous protocols ^28,29^. The fish lines *Tg(mnx1:GFP)^ml2^*, *Tg(huC:egfp)*, *Tg(huC:mcherry)* and *Tg(gfap:egfp)* have been described in the previous work ^28,29^. The whole mount *in situ* hybridization (WISH) was performed as previously described ^30^. The templates for generating antisense RNA probes were amplified from the cDNA library, using specific primers targeting *tyw1* and *gpam* listed as below:

*tyw1*-Forward: 5’-AAGGCCAGCTGCAAGAATAA-3’
*tyw1*-Reverse: 5’-GTGTGGTGTCTCCAGCAGAA-3’
*gpam*-Forward: 5’-TCCTGTTCACAAGTCCCACA-3’
*gpam*-Reverse: 5’-AGTTCTTGCGGAGCATCCTA-3’

#### Morpholino and mRNAs Injections

*tyw1* splice-blocking Morpholino (MO; Gene Tools, OR, USA) sequence was 5′- AACCTTATTCCCACTTAATGTTACC-3′. *gpam* splice-blocking Morpholino sequence was 5′- GGTGCTACTTTTCTCCAAGCTTACC-3′. The sequence of a standard control MO oligo was 5′-CCTCTTACCTCAGTTACAATTTATA-3′. *tyw1* Translation-blocking Morpholino sequence was 5’-CAGCATCTCATGTACTCTCTCCATC-3’. *gpam* Translation-blocking Morpholino sequence was 5’-ACGTCCATCCCCTCTCTTCAAACCA-3’. The MOs were diluted to 0.3 mM with RNase-free water and injected into the yolk of one to two-cell stage embryos and then raised in E3 medium at 28.5 °C. The wild-type and mutated cDNAs (*TYW1* (NM_018264) mut1: p.R389Q; *TYW1* mut2: p.R206C; *GPAM* (NM_001244949) mut1: p.G499R; *GPAM* mut2: p.P669S) containing the open reading frame of the zebrafish and human *tyw1* or *gpam* genes were cloned into pCS2+ vector respectively and then were transcribed *in vitro* by using the mMESSAGE mMACHIN Kit (Thermo Fisher Scientific, MA, USA) after the recombinant plasmids linearized with NotI Restriction Enzyme (NEB, MA, USA), and then the capped mRNAs were purified by RNeasy Mini Kit (Qiagen, Frankfurt, Germany). Around 2 nl mRNA were injected at 50 ng/μl into 1/2-cell stage embryos.

#### RNA isolation, reverse transcription, and PCR

Total RNA was extracted from zebrafish embryos by TRIzol reagent according to the manufacturer’s instructions (Invitrogen, Thermo Fisher Scientific, MA, USA). The reverse transcription was carried out according to the standard protocols from manufacturer (Fermentas, Thermo Fisher Scientific, MA, USA). The PCR was carried out as previously described to validate the efficiency of splicing-blocking effect of specific morpholino ^28^ by using the primer sequencing listed as below.

*tyw1*-Forward: 5’-TTATCGGTGTTGTCGGGTTT-3’
*tyw1*-Reverse: 5’-CCTCTGCCAATCTGTCTTCC-3’
*gpam*-Forward: 5’-ACAGTGCGCAAAAAGAGGTC-3’
*gpam*-Reverse: 5’-TAAGCACGTTCTCCACCACA-3’

#### Locomotion analysis in zebrafish larvae, microscopy and statistical analysis

The locomotion analysis of 5 dpf zebrafish larvae was carried out by using the DanioVision system (Noldus Information Technology, Wageningen, Netherlands). The confocal imaging was performed by using a TCS-SP8 LSM confocal imaging system (Leica, Wetzlar, Germany). The zebrafish embryos were embedded as we previously did ^28^. Photographs of *in situ* hybridization results were taken by using a DP70 camera on an Olympus stereomicroscope MVX10. Statistical comparisons of the data were carried out by student’s t-test, and P < 0.05 was considered statistically significant.

### Mice modeling

All mice experiments in this study were approved by the institutional animal care and use committee in the Guangzhou Medical University (registration No. 2019-436, 2019-694). In this study, all mice, either wild-type or mutant, were generated from the C57BL/6J strain and provided by the Cyagen Biosciences (Guangzhou, China).

#### Generation of knockout mice by CRISPR/Cas9 system

Knockout mouse lines were created by CRISPR/Cas9-mediated genome engineering. To create a *Tyw1* knockout mouse model, exon 2~3 was selected as the target site. gRNA was designed as the following sequences:

gRNA1 (matching reverse strand of gene): GCCTAAGGACCACGTTTCGATGG.
gRNA2 (matching reverse strand of gene): CATGTAAGACCTCACGATCAAGG.

To create a *Gpam* knockout mouse model, exon 2~4 was selected as the target site. gRNA was designed as the following sequences:

gRNA1 (matching reverse strand of gene): CTCCCACGGGAAAACCACGCAGG.
gRNA2 (matching reverse strand of gene): CCCAAGGGGTCACGCACCACAGG.

Cas9 mRNA and gRNA were co-injected into the cytoplasm of fertilized eggs. The microinjected zygotes were cultured in medium until the two-cell stage, and then were transferred into the oviducts of pseudopregnant ICR females at 0.5 d after mating with vasectomized males. The mutant F0 mice were identified by genomic PCR and the DNA sequence was confirmed by Sanger sequencing. Subsequent breeding of the mutant F0 mice generated offspring with desired genotypes for experiments.

The primers used for genotyping and gene editing evaluation were listed as below. Gene knockout efficiency was identified using qPCR (supplementary figure 16).

Primers for knockout mice models:

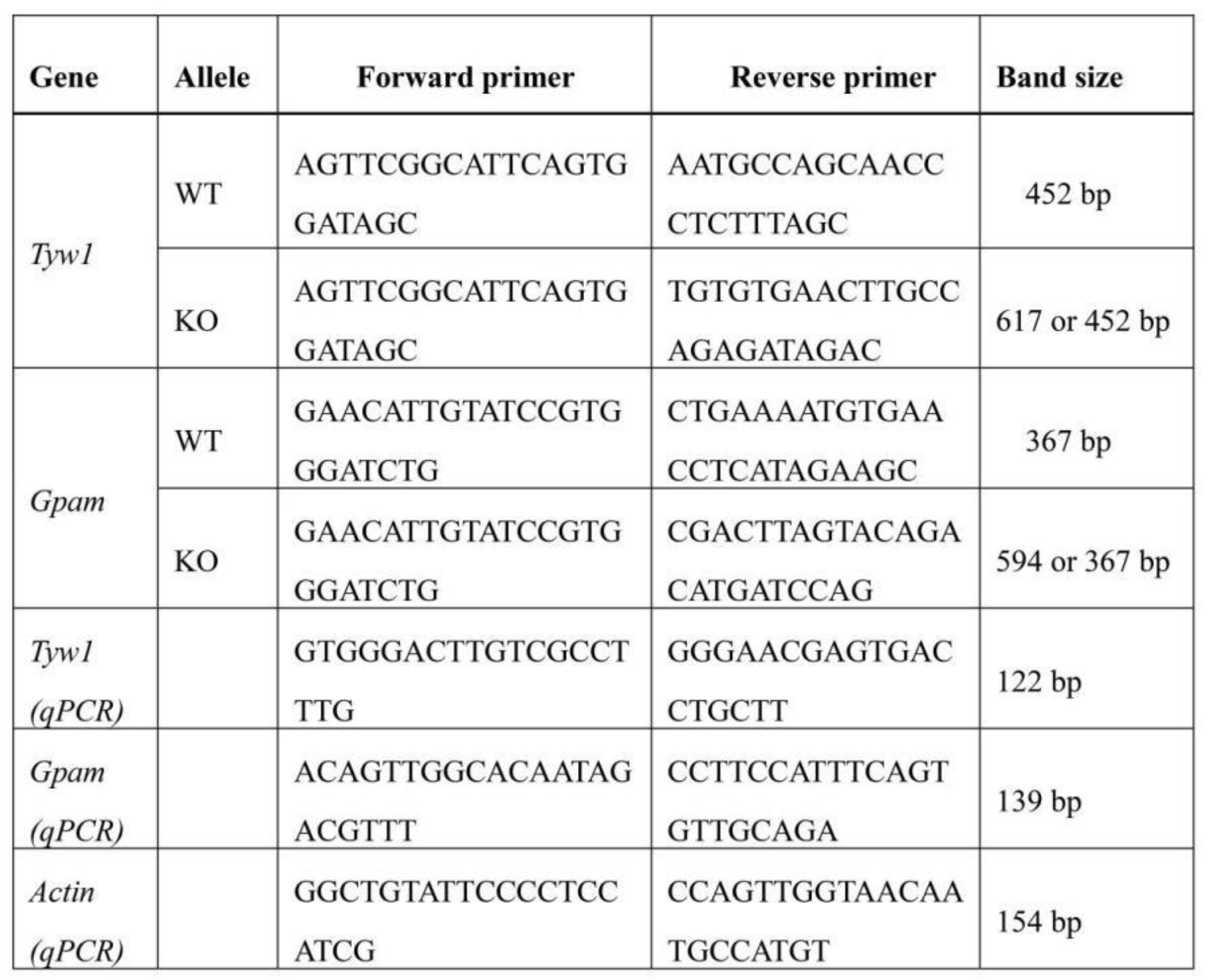

#### Behavioral tests

All behavioral tests were carried out on both male and female mice at 8 weeks of age, and the balanced gender was kept for both wild-type and *Tyw1*−/− or *Gpam*−/− mice. Each test was conducted at fixed day time (between 8:30 am to 18:30 pm) on each training day. Mice were moved to the testing room 1 h before behavioral testing for acclimation, and those participating in multiple tests were allowed to rest for at least 3 days between two tests. All experimental areas were cleaned with 70% ethanol before the tests and between subjects. All behavioral tests were carried out with the presence of two researchers blinded to the genotype.

#### Morris water maze test

Morris water maze test was performed to investigate the learning and memory ability of mice. Briefly, a platform was placed at the central zone of one quadrant of the pool below the surface of water. The mice were trained to learn the position of the hidden platform for 4 days. At day 5, the platform was removed. The mice were released into water and allowed to swim for 60 s to search a virtual quadrant centered on the location of the platform. Duration in zone of platform refers to amount of time mice remained in this virtual zone. Frequency of crossing platform refers to number of times mice crossed the virtual zone, while latency to first arrival refers to the time needed for mice to reach the virtual zone at first time, while distance travelled in target quadrant refers to the distance mice swam in the virtual quadrant. Videos were recorded and analyzed by using the SuperMaze V2.0 software (XinRuan, Shanghai, China).

#### Rotarod test

Rotarod test was performed to indicate motor coordination of mice in each group. Firstly, mice were trained three times at the rate of 30 rpm before test, and each training maintained for 5 min. In the test, the rotation rate increased gradually to reach 40 rpm within 300 s. The test was over when a mouse fell off. Each mouse was detected for 3 times and the mean latency time at rotarod was recorded.

#### Grip strength test

Grip strength of mice forelimbs was measured with a grip strength test meter (BIOSEB, EB Instruments). During the test, the mice were placed over the grid allowing only forepaws to attach to the grid. After the mice paws grasped the grid, their tails were pulled horizontally until they completely released hold. Each mouse was tested for three times. The readings of grip strength and duration of grasping the grid were recorded and analyzed by using the SuperGSM software (XinRuan, Shanghai, China).

#### Tissue preparation

Mice were anaesthetized with 50 mg/kg sodium pentobarbital. After perfused with 0.9% NaCl and 4% paraformaldehyde, brains were extracted and post-fixed for 24 h, followed with dehydration by using sucrose. Brains were cut into two hemispheres, and then serial sagittal sections (30 μm thick) were sliced with freezing microtome (CM3050S, Leica, Germany).

#### Nissl staining

Sections were immersed into 75% ethanol (30 s), dH_2_O (30 s) and cresyl violet (2 min). Then, sections were dehydrated with gradient ethanol (75%, 95%, 100%) for 30 s respectively, followed by incubation with xylene and mounted with neutral resins. The images were taken under an inverted microscope.

#### 5-Ethynyl-2’-deoxyuridine labeling

For proliferation assays, pregnant dams on E13.5 were intraperitoneally injected with 5-Ethynyl-2’-deoxyuridine (EdU, 20 mg/kg body weight) 2 h before sacrifice. Embryonic brains were harvested, fixed in paraformaldehyde and immersed in 30% sucrose for dehydration. Brain slices were then stained with Click-iT EdU Imaging Kits (C10339, Invitrogen, Thermo Fisher Scientific, USA). Cells incorporated with EdU were determined under a fluorescence microscope. Percentage of EdU^+^ fluorescence area (indicating the cell number of EdU^+^) of E13.5 mouse brain cortex was calculated in 3 random fields per coverslip. For migration assays, pregnant dams on E15.5 were intraperitoneally injected with EdU. Brains were harvested after 72 h and stained as above.

#### Immunostaining

To detect expression of TYW1 and GPAM in human brain tissue, the normal brain tissue microarrays (BNC17011, Biomax, MD, USA) were prepared. According to the instruction from Biomax official website, BNC17011 were derived from normal brain tissue of male or female at the age of 2 to 50, including brain regions such as frontal lobe, apical lobe, occipital lobe, temporal lobe, midbrain, pons, medulla oblongata, thalamus opticus, cerebellum, hippocampus, callositas, optic nerve, and spinal cord. The slides were immersed in xylene and graded ethanol to deparaffinize and rehydrate. After antigen retrieval using microwave with sodium citrate buffer, 0.3% Triton-X-100 was added to permeabilize the tissue. The slides were blocked with goat serum and incubated with primary antibodies at 4 °C overnight. After washing with PBS, the sections were incubated with secondary antibodies.

For mouse brain tissues, the perfused brain samples were fixed within 4% paraformaldehyde (PFA) in 0.01 M PBS at 4 °C for 24 h, then washed by PBS twice, cryoprotected with sucrose gradients, snap frozen, and sectioned with a cryostat (Leica, Wetzlar, Germany) to thickness of 30 μm. Then the sections were immuno-stained by using aforementioned methods, except deparaffinization and rehydration.

Number of Cux1^+^ or Foxp2^+^ cells in frontal cortex and motor cortex were counted on three sagittal cerebrum sections, respectively. Gfap^+^ and NeuN^+^ cells in different brain regions were counted in 3 random fields per section. Data were obtained from at least three independent experiments and analyzed with ImageJ software.

In order to visualize and measure the cholesterol level, Filipin staining was performed in tissue sections (10μm thick) by using a cell-based Cholesterol Assay Kit (ab133116, Abcam, UK). Briefly, the tissue sections were washed (3 × 5 min) with Cholesterol Detection wash buffer, and then the Filipin III was added to each section and incubated in the dark for 60 min at room temperature. After washing, the fluorescence images were obtained by a Leica DMi8 (Leica Microsystems, Wetzlar, Germany) fluorescence microscope by using × 20 HC PL FLUOTAR objective.

#### Western blot analysis

Total protein was extracted from peripheral blood using cell lysate containing RIPA and protease inhibitor cocktail. And the brain tissues were lysed with 2% SDS in PBS with PMSF and proteinase inhibitor cocktail. The BCA assay was used to determine protein concentration. Proteins were resolved on 7.5%, 10% or 15% tris-glycine gels based on different molecular weight and transferred to PVDF membranes. After blocking, the membranes were incubated with primary antibodies, followed by horseradish peroxidase (HRP)-labeled secondary antibodies. Then, blots were visualized by West Pico Plus Chemiluminescent Substrate (Thermo Fisher Scientific, MA, USA) and scanned using ChemiDoc™ MP system (Bio-Rad, CA, USA). Densitometries of individual blot signals were quantified using ImageJ software.

#### qRT-PCR analysis

Total RNA was extracted from the mice brain tissues by using TRIZOL reagent (Invitrogen, Thermo Fisher Scientific, MA, USA), and cDNA was synthesized from 1 μg total RNA by using PrimeScript™ RT Master Mix (RR036Q, Takara, Shiga, Japan). Quantitative PCR was carried out by using PowerUp™ SYBR™ Green Master Mix (A25778, Applied Biosystems, Thermo Fisher Scientific, MA, USA) on Biosystems QuantStudio 6 Flex Real-time PCR system (Applied Biosystems, Thermo Fisher Scientific, MA, USA). *β -Actin* was used as a reference gene. For comparison, mRNA expression level of knockout mice were normalized to those of wild-type mice for each subject. The primers were:

*Tyw1*-Forward: 5’-GTGGGACTTGTCGCCTTTG-3’
*Tyw1*-Reverse : 5’- GGGAACGAGTGACCTGCTT-3’
*Stil*-Forward: 5’- GACACAATTCAGGACTGGTAGAC-3’
*Stil*-Reverse : 5’- GGCATGATCCACTTTCTGTTCA-3’
*Sass6*-Forward: 5’- ATTCCTTTACGCGGACTTAGC-3’
*Sass6*-Reverse : 5’- AAGTAGGCTGAAGACGAGGAG-3’
*Ncapd2*-Forward: 5’- AGCCAGACAAGCCTCATTGAC-3’
*Ncapd2*-Reverse: 5’- TCCATAGGTGACGGATGTCCA-3’
*Cenpe*-Forward: 5’- CTTCAGTGGCTGTCTGTGTTC-3’
*Cenpe*-Reverse: 5’- CCATCGCTCTGATAAATAGCGTT-3’
*β -Actin* -Forward: 5’- GGCTGTATTCCCCTCCATCG-3’
*β -Actin* -Reverse: 5’- CCAGTTGGTAACAATGCCATGT-3’

#### Culture of primary neurons

E13.5 mouse embryo brains were taken out from *Tyw1*+/− pregnant dam. Cortex in each brain was dissected separately and collected in Hibernate-E supplemented with 2% B27 on ice. Single cells were obtained by using 0.05% trypsin (containing 0.2 mM EDTA) digestion for 10 min at 37 °C. After filtration with 70 μm strainer and centrifugation, cells resuspended in Neurobasal medium with 2% B27 Plus Supplement, 0.25% Glutamax and 25 μM glutamate were placed in poly-D-lysine coated plates. Adherent neurons were prepared for EdU and TUNEL staining. To prepare migration assay, isolated cells were initially cultured in ultra-low attachment plates to form neurospheres, which could be digested by using 0.05% trypsin without damage of neurites.

#### Proliferation and apoptosis assays of primary neurons

At 2 DIV, cells were labeled with 10 μM EdU solution and incubated for 4 h. Cells were treated by using 4% PFA, followed by 0.5% TritonX-100 permeabilization. After being washed twice with 3% BSA, cells were incubated with Click-iT reaction cocktail according to protocol of Click-iT EdU Imaging Kits (C10339, Invitrogen, Thermo Fisher Scientific, MA, USA) and stained with DAPI. Dead cells were labeled using *In Situ* Cell Death Detection Kit, POD (11684817910, Roche, Switzerland).

#### Migration test of primary neurons

Neurospheres were digested with 0.05% trypsin to obtain single neurons and resuspended in Neurobasal Medium with 1% B27 Plus Supplement. Cells were seeded on the upper layer of a cell culture insert with PET track-etched membrane (8 μm pore size, 353097, Corning, NY, USA) at density of 1 × 10^5^ cells/well. Neurobasal Medium with 2% B27 Plus Supplement was added into the bottom of the lower chamber. After 16 h, the culture insert was taken out and the medium was removed carefully. Cells on the upper surface were wiped, while migrating cells in pore and on the lower surface of membrane were fixed, stained with 0.1% crystal violet and observed.

#### Flow cytometry

Whole brain tissues from *Gpam*−/− mice on P1 (postnatal day 1) were collected, including brain tissues of their wild-type littermate. After digestion with 4 mg/ml papain and 0.1 mg/ml DNase in 37 °C shaker for 30 min, suspension was diluted using DMEM/F12 medium containing 5% FBS and filtered with 70 μm strainers. Then, suspension was centrifuged and the cells were fixed in 4% PFA. 0.3% Triton-X100 was used for permeabilization. After being blocked with 5% goat serum, cells were incubated with mouse anti-Gfap antibody and rabbit anti-Ki67 antibody or rabbit anti-Caspase-3 antibody at 4 °C for 1 h. Cells were washed with DPBS and incubated with Alexa Fluor 488 goat anti-mouse IgG and Alexa Fluor 647 goat anti-rabbit IgG at 4 °C for 30 min. After being washed for two times, cells were suspended by using DPBS and analyzed on the flow cytometer (BD FACSCanto, BD Biosciences, CA, USA).

#### Measurement of phosphatidic acid (PA)

Phosphatidic acid was measured by using the PicoProbe™ Phosphatidic Acid Assay Kit (BioVision, CA, USA). In brief, medulla of wild-type and *Gpam*−/− mice was homogenized in PA assay buffer. Lipid extraction was obtained according to the protocol and solubilized in 5% Triton X-100 solution. “Sample background control” and “Sample” were prepared in parallel. Standard curve was generated by using PA standard solution. Converter mix was only added in sample and standard wells. All wells were incubated at 45 °C for 1 h. Reaction mix was added in each well and incubate at 37 °C for 30 min. Fluorescence was recorded at Ex/Em = 535/587 nm and PA concentration was expressed as nmol PA per mg tissue weight. Value of PA concentration was normalized.

#### Measurement of phosphatidylcholine (PC)

Phosphatidylcholine was measured by using the phosphatidylcholine assay kit (Abcam, UK). For each individual, about 10 mg brain tissues were washed with cold PBS, resuspended in the assay buffer, and homogenized on ice. After 10 min incubation, samples were centrifuged for 5 min at 4 °C at 16,000 g. The supernatant was incubated with the reaction mix including OxiRed Probe supplemented with hydrolysis enzyme for 30 min. The colorimetric reading was measured at OD_570_ nm on a microplate reader.

#### Measurement of Phosphatidylethanolamine (PE)

Phosphatidylethanolamine was measured by using the phosphatidylethanolamine assay kit (Abcam, UK) according to the manufacturer’s instruction. Briefly, solubilized lipids were extracted from brain tissues with 5% Triton X-100, incubated with converter mix at 45°C for 1 hour, and then with reaction mix at 40 °C for 3 hours. Fluorescence was recorded at Ex/Em 535/587 nm.

### Modeling the effect of *TYW1* knockout on ribosomal frameshift and the calculation of attenuation coefficient for proteins

The modeling of *TYW1*-knockout effect on ribosomal frameshift was based upon the fact that TYW1 is a critical enzyme involved in the synthesis of wybutosine, so the hypomorphic and null alleles of *TYW1* could reduce and even remove the production of wybutosine accordingly ^31,32^. It is known that wybutosine at the 37 position of tRNA^Phe^ near the 3’ base of anticodon can stabilize the codon-anticodon interaction thus drastically reduce the ribosomal frameshift upon a specific codon UUU ^33^. We, therefore, hypothesized that the density and distribution of codon UUU along a specific mRNA should determine the extent of influence exerted by wybutosine, and in parallel, the relative abundance of wybutosine was another major effector during this process. Our modeling procedure went as follows:

1. Download the UCSC RefSeq (refGene) gene models of human, mouse, and zebrafish from https://genome.ucsc.edu/cgi-bin/hgTables. Extract the coding domain sequences (CDS) of each protein-coding gene. If there exists multiple isoforms, choose the longest ones.
2. Label the location of codon UUU, normalize the location values by the length of CDS. By doing this, we harvested a series of normalized location values, i.e., L(1), L(2), …, for each gene, where 0 < L(n) < 1, n = 1, 2, ‥‥
3. We made the following two assumptions: (1) for a codon UUU at any location, there is a fixed chance of ribosomal frameshift designated by Pf, given a fixed level of wybutosine; (2) for a given codon UUU located at L(n), after frameshifting, the remaining activity of a protein is proportionate to L(n). Thus, the consequence of translating a mRNA holding multiple codon UUU, i.e., the remaining activity of a protein, designated by R, can be estimated as:

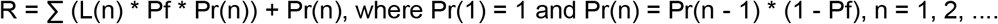
4. For the wild-type (WT) and *TYW1*-knockout mice (KO), we defined the R values for WT and KO to be R(WT) and R(KO), respectively. The R value is determined by the density and distribution of codon UUU in a given mRNA, also by Pf (the chance of ribosomal frameshift at a specific location of mRNA, given a fixed level of wybutosine). We set the values of Pf for R(WT) and R(KO) to be 0.12 and 0.35, respectively ^33^. Subsequently, we defined the **attenuation coefficient** to be R(KO) / R(WT). An attenuation coefficient reflected the extent of protein activity reduction with *TYW1*-knockout, and the expected values of attenuation coefficient ranged from 0 to 1. The lower the value of attenuation coefficient, the more severe the ribosomal frameshift and protein degradation when *TYW1* is knocked out.
5. After generating a list of attenuation coefficients for each human / mouse / zebrafish protein, we focused on proteins meeting with these criteria: (1) a protein with homologues in all three species; (2) a protein with attenuation coefficient less than 0.34 (i.e., 0.12 / 0.35); (3) a protein with its corresponding gene matching *TYW1* in terms of expression profiles in normal brains; (4) a protein known to be disease-related, with records in HGMD or OMIM.

### Reagents

| Reagent type | Designation | Source | Identifiers | Dilution |
| --- | --- | --- | --- | --- |
| Antibody | Self-generated anti-TYW1<br>(rabbit polyclonal) | GenScript | Cat no: 2469 | 1:20 |
| Antibody | anti-GPAM<br>(mouse monoclonal) | Santa Cruz | Cat no:sc-398135 | 1:500 |
| Antibody | anti-NeuN<br>(mouse monoclonal) | Millipore | Cat no: MAB377B<br>RRID: AB_177621 | 1:150 |
| Antibody | anti-GFAP<br>(mouse monoclonal) | Millipore | Cat no: MAB360<br>RRID: AB_11212597 | 1:400 |
| Antibody | anti-CUX1<br>(mouse monoclonal) | Abcam | Cat no: ab54583<br>RRID: AB_941209 | 1:40 |
| Antibody | anti-FOXP2<br>(rabbit polyclonal) | Abcam | Cat no: ab16046<br>RRID: AB_2107107 | 1:1000 |
| Antibody | anti-MAP2, clone AP20<br>(mouse monoclonal) | Millipore | Cat no: MAB3418<br>RRID: AB_94856 | 1:100 |
| Antibody | anti-MBP<br>(rat monoclonal) | Abcam | Cat no: ab7349<br>RRID: AB_305869 | 1:50 |
| Antibody | anti-CNPase<br>(mouse monoclonal) | Millipore | Cat no: MAB326<br>RRID: AB_2082608 | 1:200 |
| Antibody | anti-Ki-67<br>(rabbit monoclonal) | Cell Signaling<br>Technology | Cat no: 9129<br>RRID: AB_2687446 | 1:400 |
| Antibody | anti-Cleaved Caspase-3<br>(rabbit monoclonal) | Cell Signaling<br>Technology | Cat no: 9664<br>RRID: AB_2070042 | 1:800 |
| Antibody | anti-SAS6<br>(mouse monoclonal) | Santa Cruz<br>Biotechnology | Cat no: sc-81431<br>RRID: AB_1128357 | 1:200 |
| Antibody | anti-CENP-E<br>(mouse monoclonal) | Santa Cruz<br>Biotechnology | Cat no: sc-376685<br>RRID: AB_11150652 | 1:100 |
| Antibody | anti-STIL<br>(rabbit polyclonal) | Abcam | Cat no: ab89314<br>RRID: AB_2197878 | 1:2000 |
| Antibody | anti-NCAPD2<br>(rabbit polyclonal) | Proteintech | Cat no: 13382-1-AP<br>RRID: AB_2149840 | 1:600 |
| Antibody | anti-β-actin (D6A8)<br>(rabbit monoclonal) | Cell Signaling<br>Technology | Cat no: 8457<br>RRID: AB_10950489 | 1:1000 |
| Antibody | Goat anti-Rabbit IgG (H+L) Highly<br>Cross-Adsorbed Secondary<br>Antibody, Alexa Fluor 568 | Thermo Fisher<br>Scientific | Cat no: A-11036<br>RRID: AB_10563566 | 1:600 |
| Antibody | Goat anti-Rabbit IgG (H+L) Highly<br>Cross-Adsorbed Secondary<br>Antibody, Alexa Fluor 488 | Thermo Fisher<br>Scientific | Cat no: A-11034<br>RRID: AB_2576217 | 1:600 |
| Antibody | Goat anti-Rat IgG (H+L)<br>Cross-Adsorbed Secondary<br>Antibody, Alexa Fluor 488 | Thermo Fisher<br>Scientific | A11006,<br>RRID:AB_2534074 | 1:300 |
| Antibody | Goat anti-Mouse IgG (H+L) Highly<br>Cross-Adsorbed Secondary<br>Antibody, Alexa Fluor 568 | Thermo Fisher<br>Scientific | Cat no: A-11031<br>RRID: AB_144696 | 1:600 |
| Antibody | Goat anti-Mouse IgG (H+L)<br>Cross-Adsorbed Secondary<br>Antibody, Alexa Fluor 488 | Thermo Fisher<br>Scientific | Cat no: A-11001<br>RRID: AB_2534069 | 1:200 |
| Antibody | Anti-rabbit IgG, HRP-linked Antibody | Cell signaling<br>technology | Cat no: 7074<br>RRID: AB_2099233 | 1:4000 |
| Antibody | Anti-rat IgG, HRP-linked Antibody | Cell signaling<br>technology | Cat no: 7077<br>RRID: AB_10694715 | 1:1500 |
| Antibody | Anti-mouse IgG, HRP-linked<br>Antibody | Cell signaling<br>technology | Cat no: 7076<br>RRID: AB_330924 | 1:4000 |
| Nuclei staining | 4',6-Diamidino-2-phenylindole<br>(DAPI) | Sigma-Aldrich | Cat no: 23670 | 1:5000 |
| Other reagent | cell-based Cholesterol Assay Kit | Abcam | Cat no: ab133116 | 1:1000 |
| Other reagent | Phosphatidylcholine Assay kit<br>(Colorimetric/Fluorometric) | Abcam | Cat no: ab83377 |  |
| Other reagent | Phosphatidylethanolamine Assay kit | Abcam | Cat no: ab241005 |  |
| Other reagent | PicoProbe™ Phosphatidic Acid<br>Assay Kit (Fluorometric) | Biovision | Cat no: K748-100 |  |

