## Supplementary figures for "In-depth analysis reveals complex molecular etiology of idiopathic cerebral palsy"

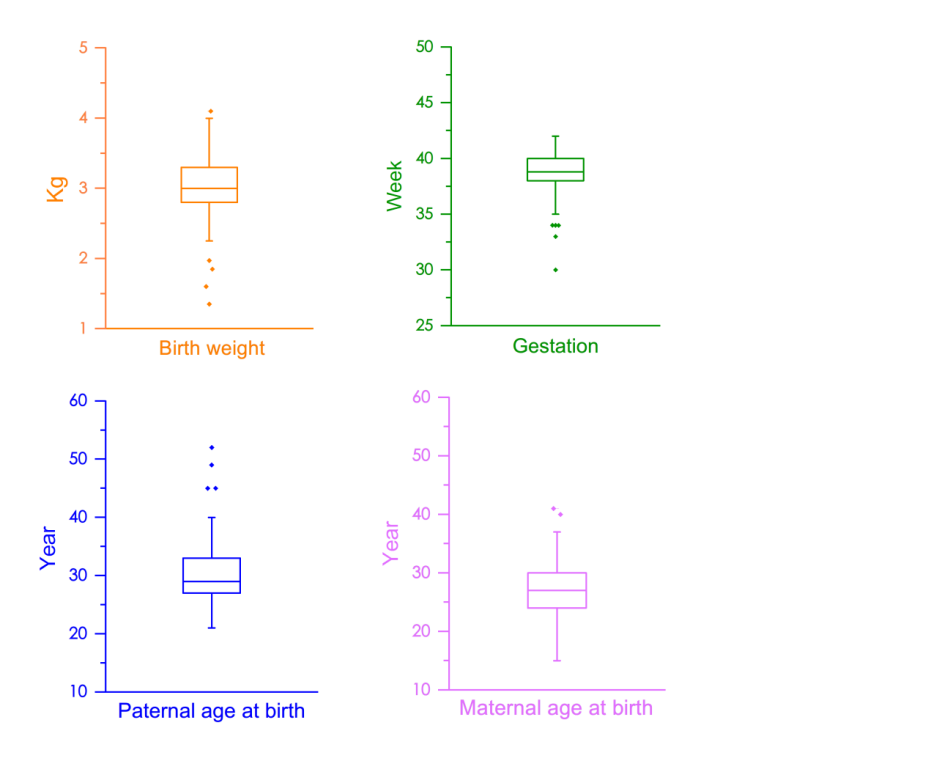

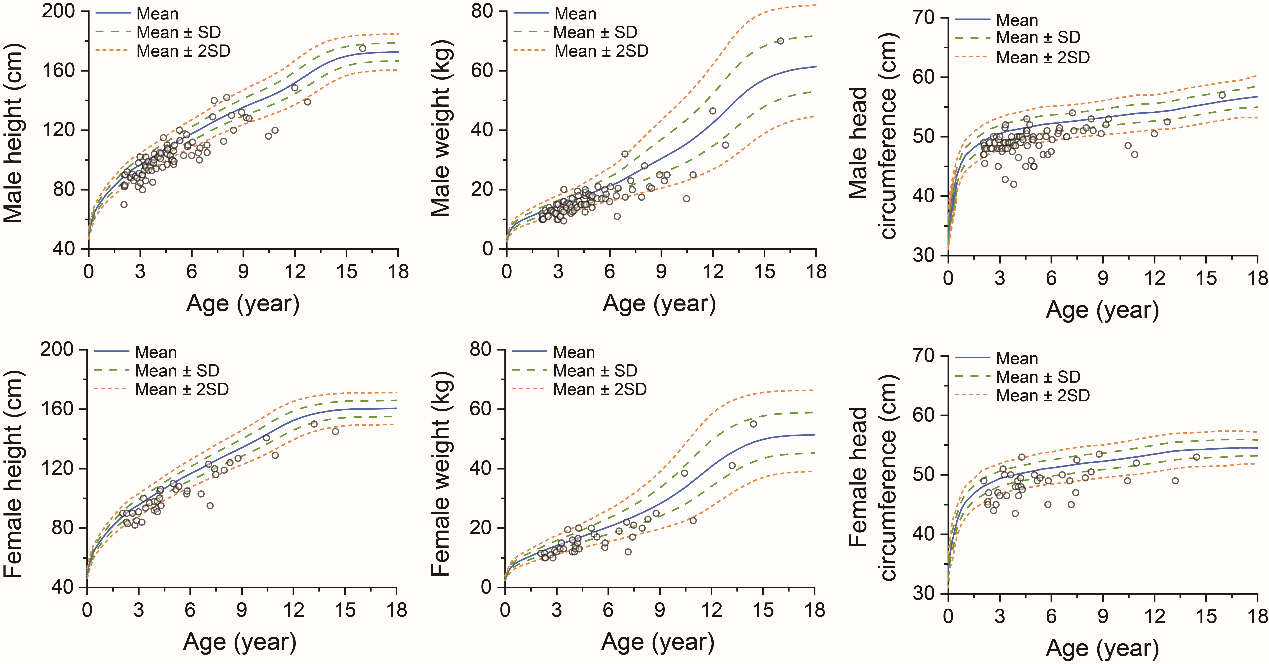

**Supplementary figure 1**: CP cohort birth information and developmental curves. ^1,2^

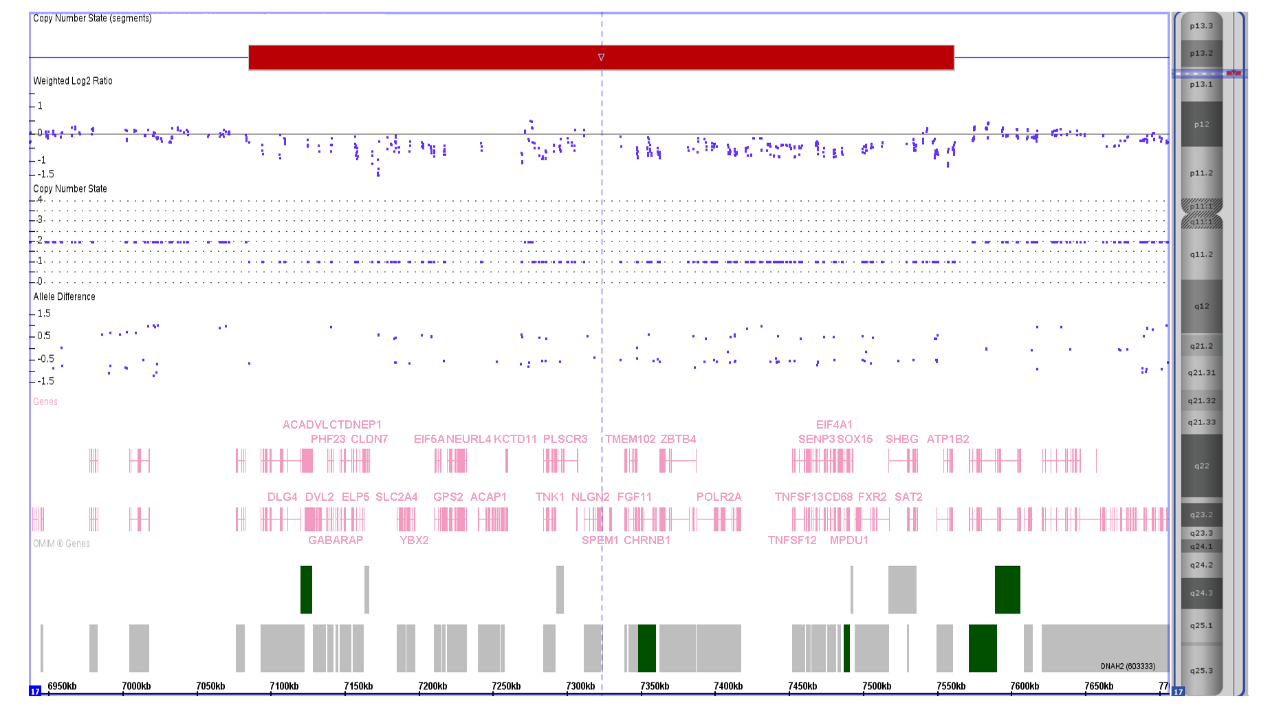

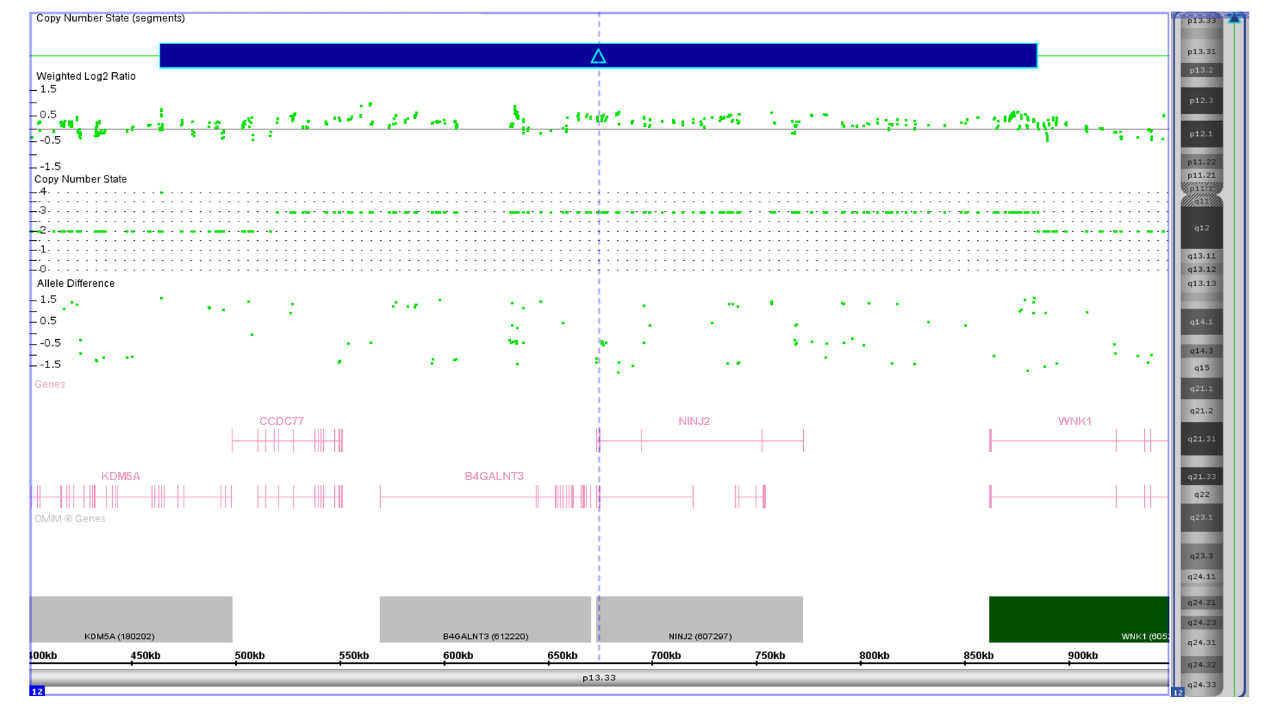

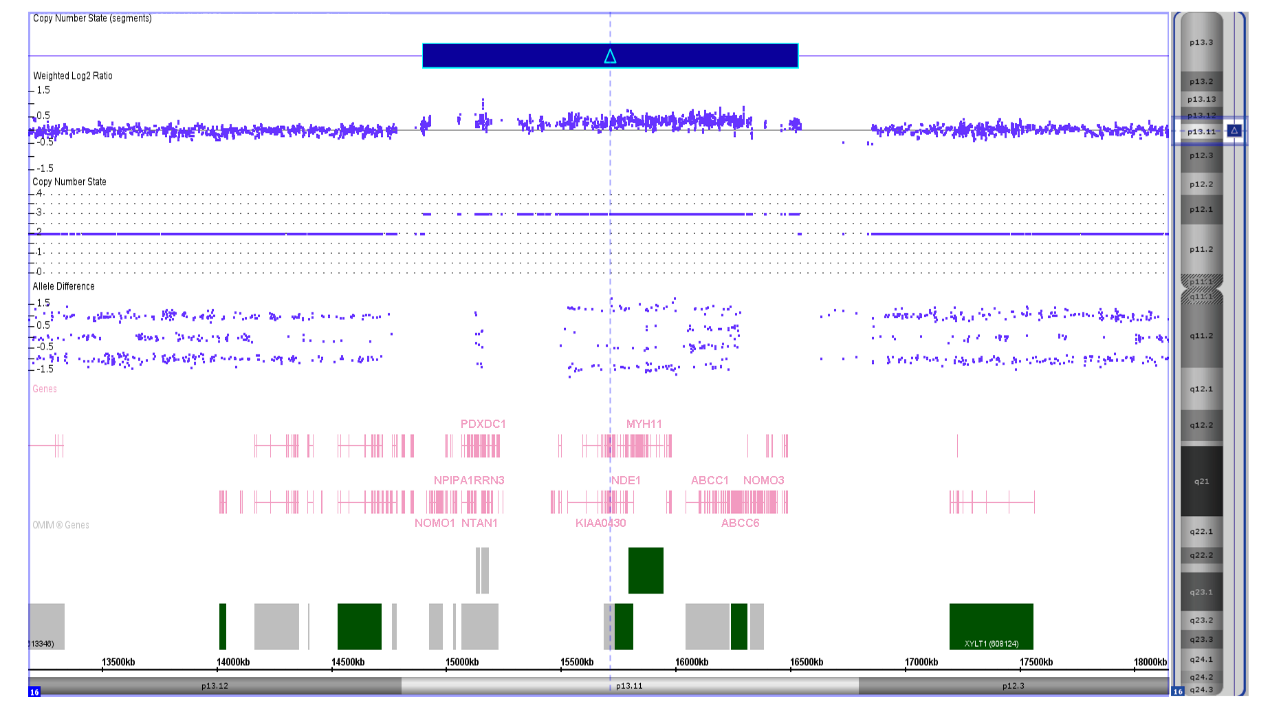

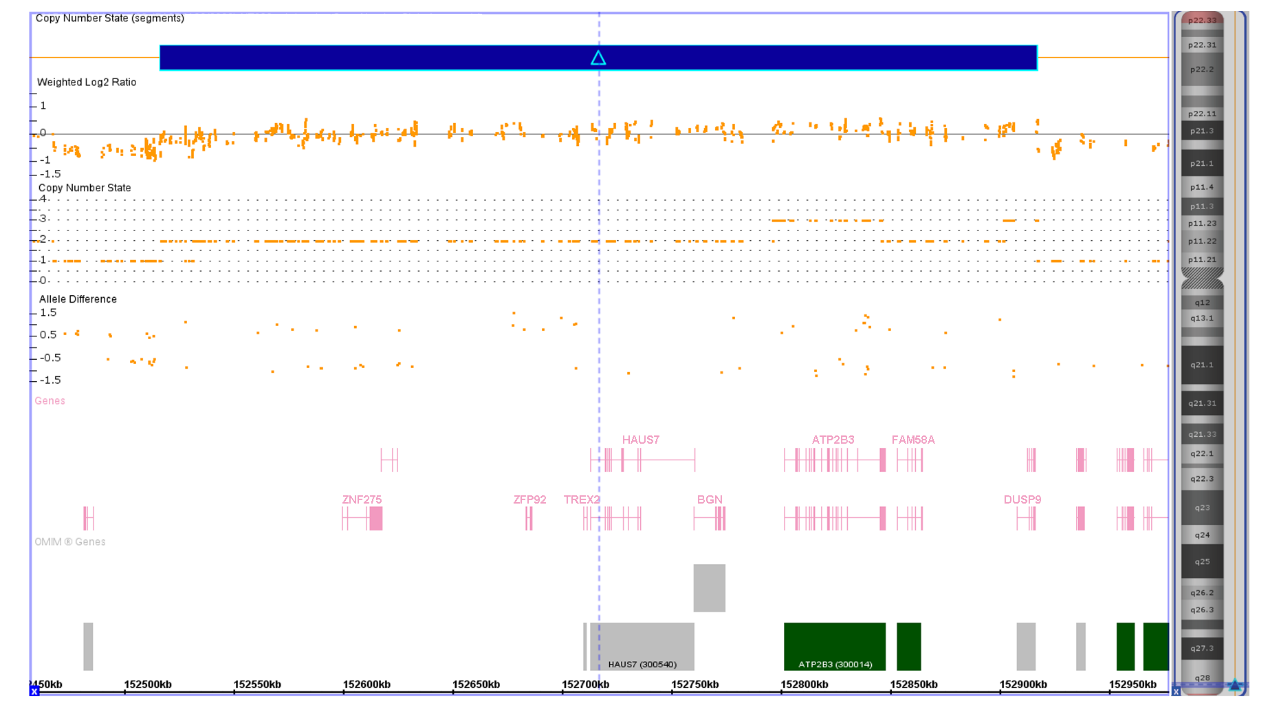
**Supplementary figure 2:** Disease-related CNVs observed in the CP cohort.

From top to bottom: arr[hg19]17p13.1(7,085,366-7,561,445)x1 in the patient CP_011_1, arr[hg19]12p13.33(464,124-885,041)x3 in the patient CP_046_1, arr[hg19]16p13.11(14,900,072-16,533,890)x3 in the patient CP_086_1, arr[hg19]Xq28(152,516,781-152,917,379)x2 in the patient CP_119_1 (male).

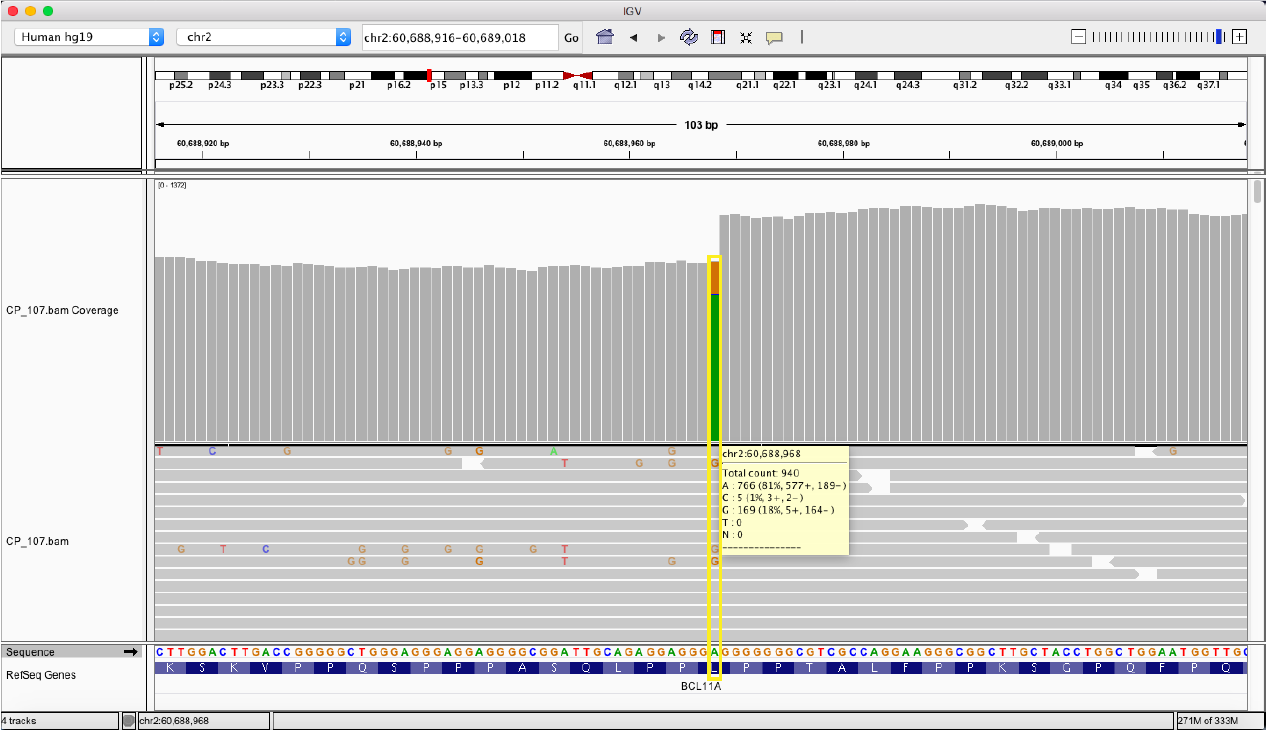

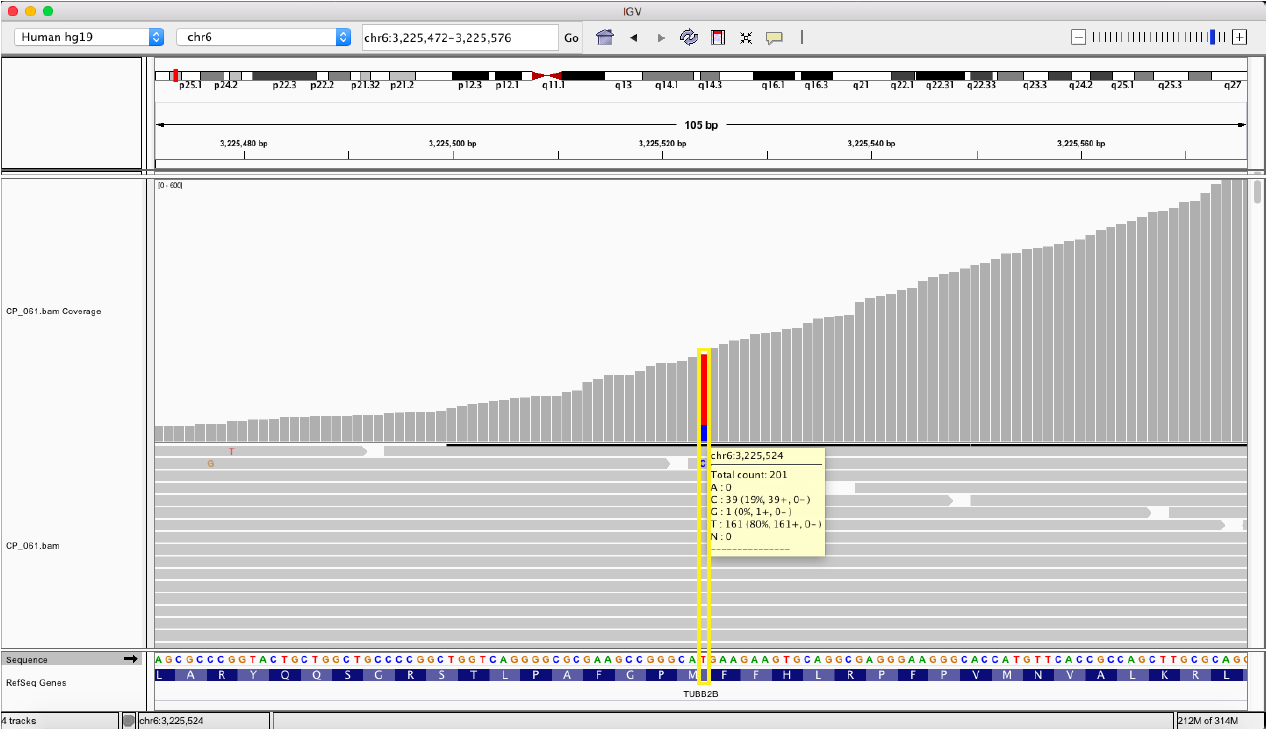

**Supplementary figure 3:** Post-zygotic mutations detected by whole-exome sequencing (600X).

Above: a A>G mutation at the gene of *BCL11A* of the patient CP_107_1 (total counts 940 of which 81% are A and 18% are G).

Below: a T>C mutation at the gene of *TUBB2B* of the patient CP_061_1 (total counts 201 of which 80% are T and 19% are C).

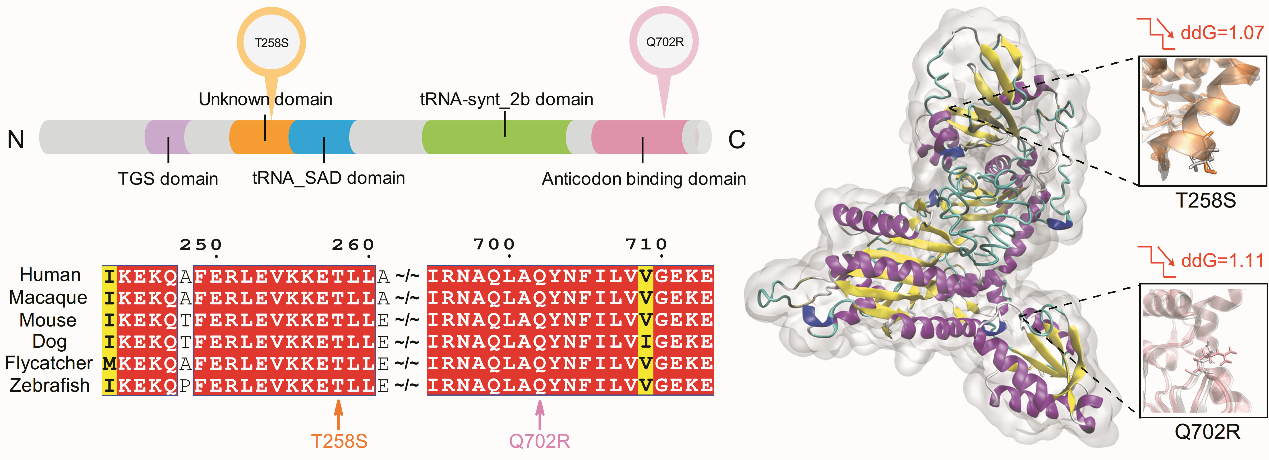
**Supplementary figure 4**: TARS and its disease-related changes. The two changes, i.e., T258S and Q702R are located in functional domains and damage the conserved sequences in different species. Both changes increase the Gibbs free-energy (ddG) thus decrease the stability of protein.

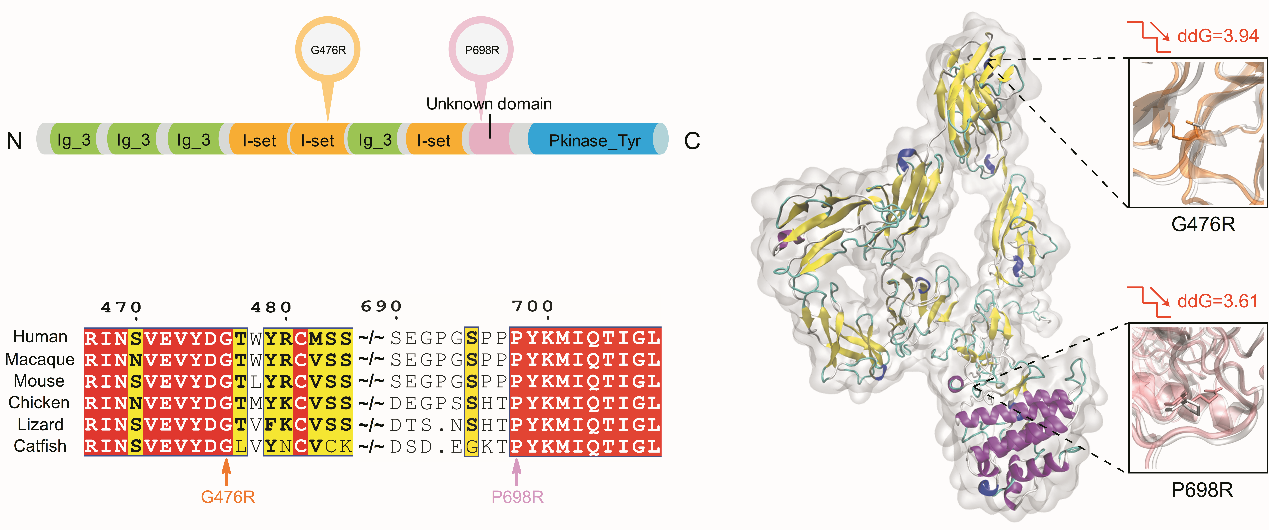
**Supplementary figure 5:** PTK7 and its disease-related changes. The two changes, i.e., G476R and P698R are located in functional domains and damage the conserved sequences in different species. Both changes increase the Gibbs free-energy (ddG) thus decrease the stability of protein.

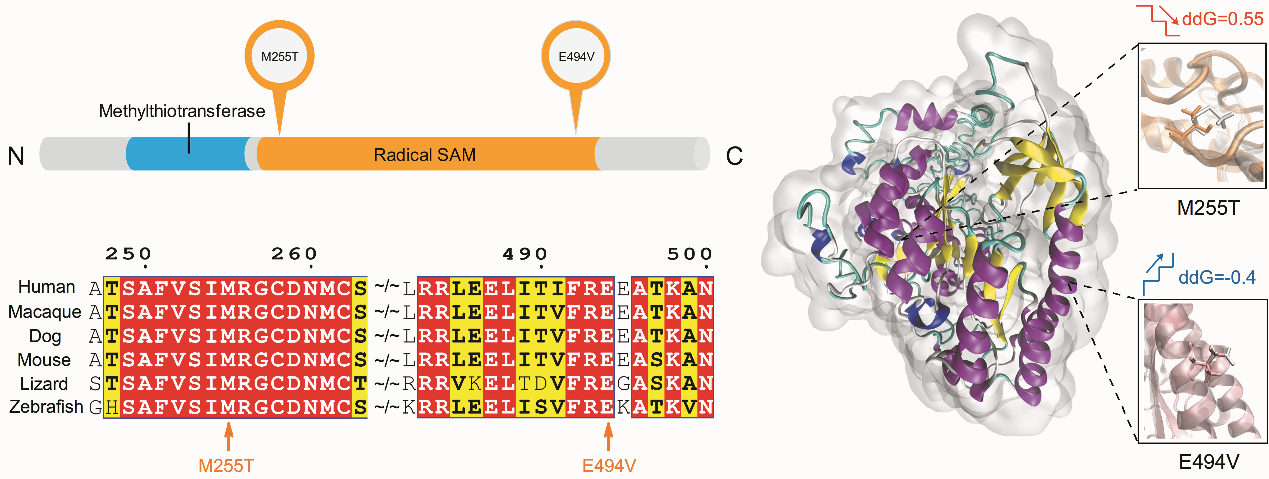
**Supplementary figure 6:** CDK5RAP1 and its disease-related changes. The two changes, i.e., M255T and E494V are located in a functional domain and damage the conserved sequences in different species. M255T increases the Gibbs free-energy (ddG) thus decreases the stability of protein.

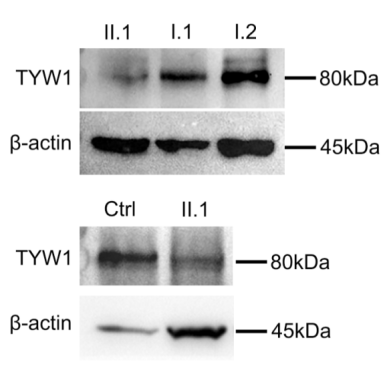

**Supplementary figure 7:** Western blot revealed significantly reduced TYW1 protein level in the patient (II.1) of family CP_012, compared to the healthy parents (I.1 and I.2) and the 5 healthy children (Ctrl) with the same age and gender.

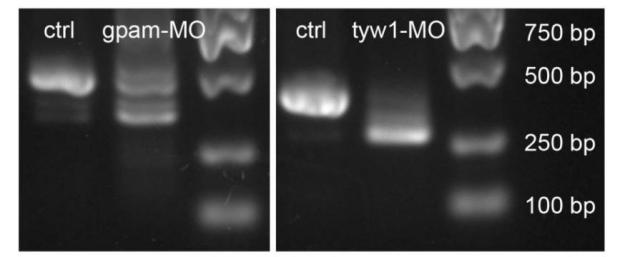

**Supplementary figure 8:** Morpholino knockdown (splice-blocking) efficiency was validated by RT-PCR in *tyw1*-MO1 and *gpam*-MO1 zebrafish.

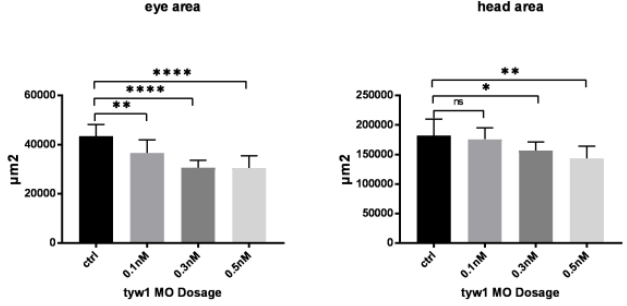

**Supplementary figure 9:** The dosage curves of *tyw1*-MO1 (a splice-blocking Morpholino) knockdown efficiency were plotted by measuring the eye area and head area of zebrafish larvae at 5 dpf. N = 10. * p < 0.1, ** p < 0.01, **** p < 0.0001, n.s.: no significant difference, unpaired t-test. dpf: days post fertilization.

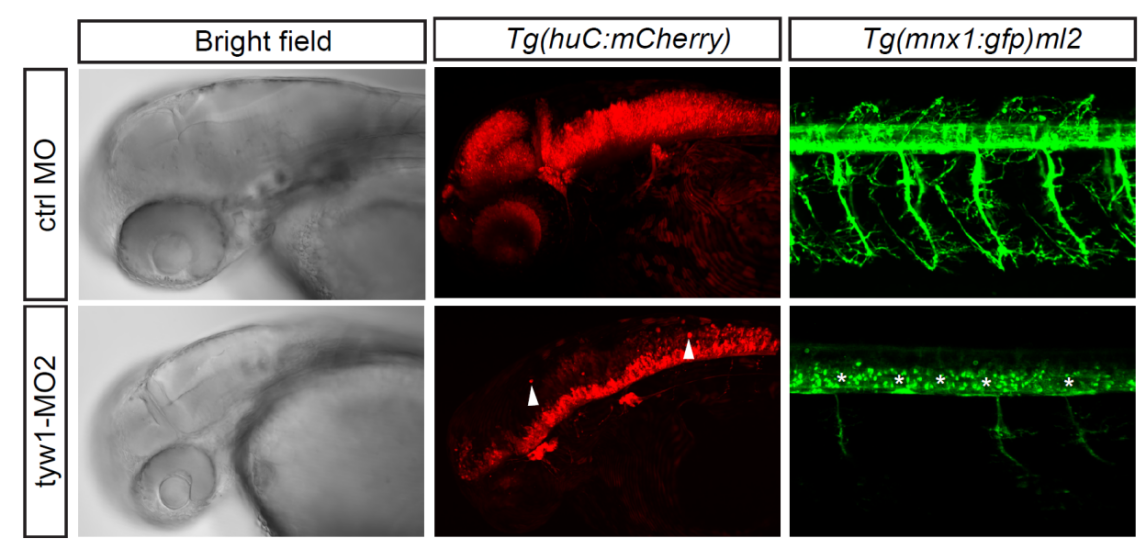

**Supplementary figure 10:** Confocal imaging of mCherry positive cells in the transgenic zebrafish *Tg(huC:mCherry)* embryos at 48 hpf showed ecotopic neuronal cells (pointed by white arrowheads) in the *tyw1*-MO2 zebrafish group by using *tyw1*-MO2 (a translation-blocking Morpholino). Confocal imaging of gfp^+^ cells in the transgenic zebrafish *Tg(mnx1:gfp)^ml2^* embryos at 72 hpf showed undifferentiated motor neuronal cells (indicated by white stars) in the *tyw1*-MO2 zebrafish group. hpf: hours post fertilization.

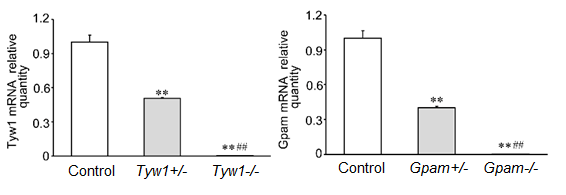

**Supplementary figure 11:** CRISPR/Cas9-mediated knockout efficiency of *Tyw1* and *Gpam* in mice. *Tyw1* and *Gpam* mRNA quantities were measured by RT-qPCR. Heterozygous knockout reduced expression of *Tyw1* and *Gpam* by about half, while homozygous knockout eliminated almost all expression of *Tyw1* and *Gpam*. N = 3. ** p < 0.01 when +/- or -/- compared with controls, ^##^ p < 0.01 when -/- compared with +/-, one-way ANOVA.

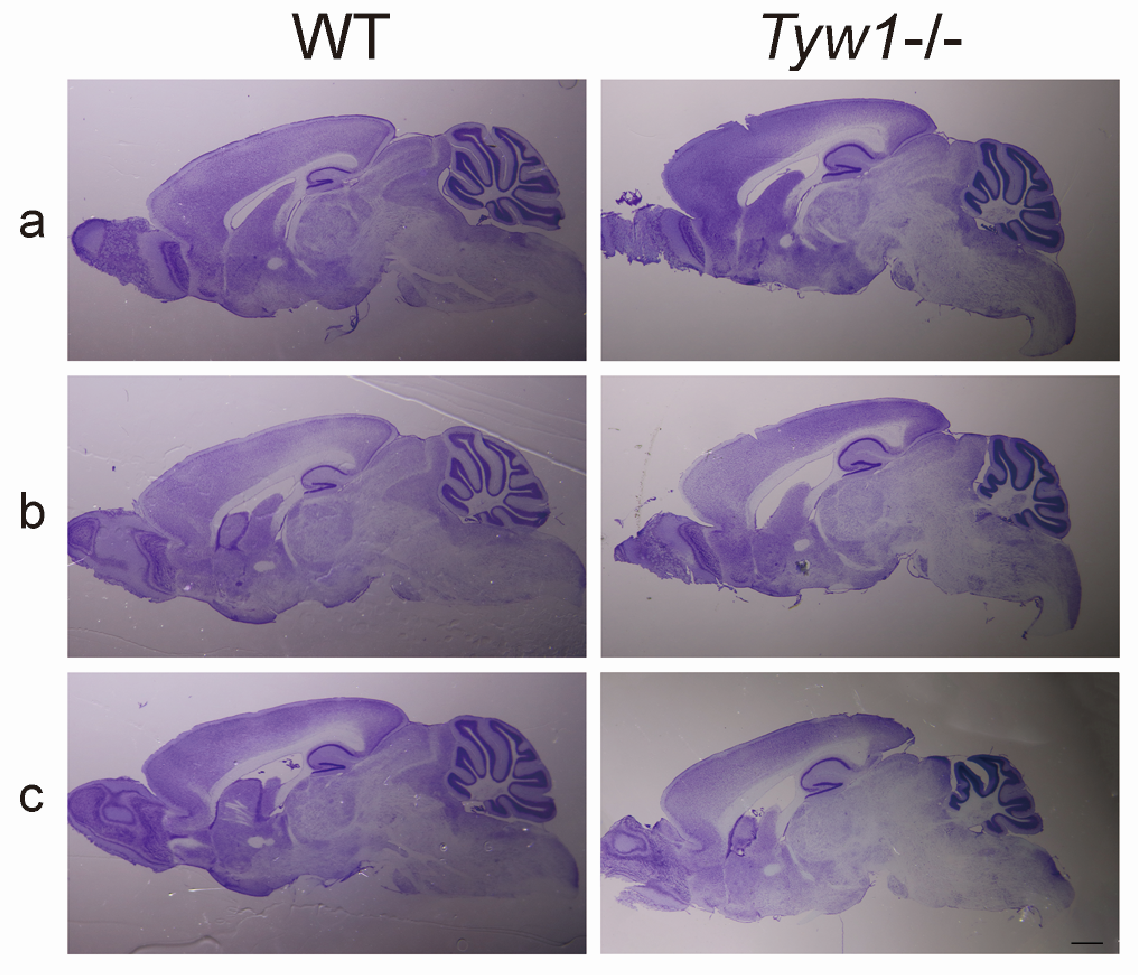

**Supplementary figure 12:** Sagittal sections of the *Tyw1-/-* and wild-type (WT) mice brains showed underdeveloped caudoputamens and enlarged ventricles among other alterations. Nissl staining, 30 μm thick. Bar = 1 mm.

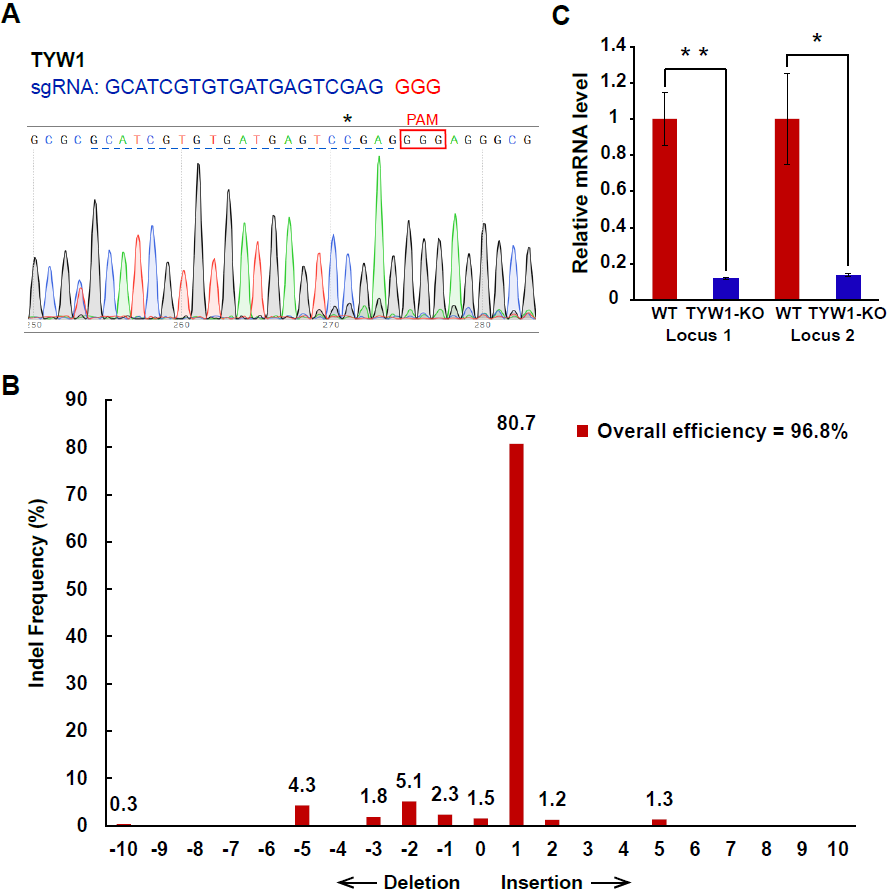

**Supplementary figure 13:** CRISPR/Cas9-mediated *TYW1* knockout in SH-SY5Y cells. **(A)** Sanger-sequencing of genome editing locus in the *TYW1*-KO SH-SY5Y cells. The sgRNA and PAM sequences were labeled by blue and red, respectively. The most frequent editing event (+1 insertion) was marked by asterisk. **(B)** Indel frequencies of *TYW1* after CRISPR/Cas9 mediated gene editing in the SH-SY5Y cells. **(C)** RT-qPCR quantification of *TYW1* mRNA level in the wildtype (WT) and the *TYW1*-KO SH-SY5Y cells. ACTB was used as internal control. The data were shown as mean ± S.D. N = 3. * p < 0.05, ** p < 0.01, unpaired t-test.

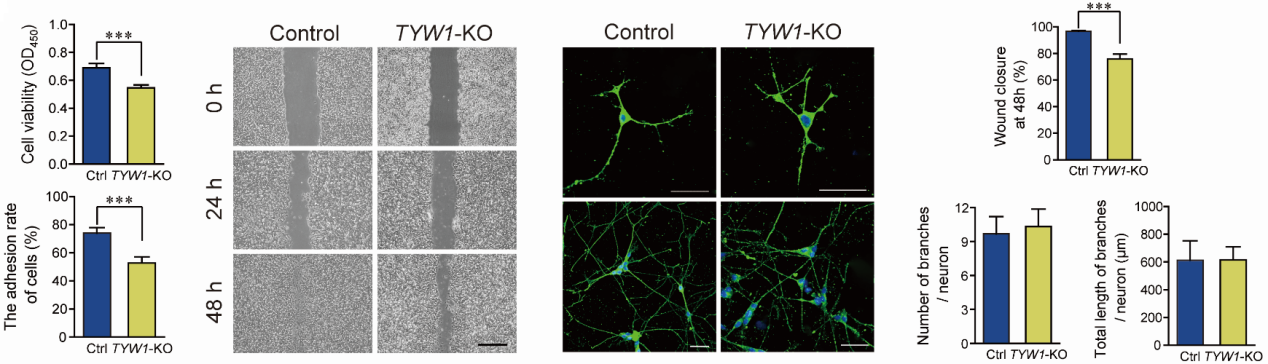

**Supplementary figure 14:** SH-SY5Y cell viability (CCK-8 assay), cell adhesion capacity (cells adhesion rate after washing), and cell migration (wound healing assay, bar = 500 μm) revealed significant decreases in the *TYW1*-KO cells versus the wild-type ones. *** p < 0.001, unpaired t-test. SH-SY5Y cell differentiation was detected after induction of RA and BDNF, and the typical arborization of MAP2-immunostained cells was observed at day 10. Bar = 50 μm. There was no significant difference between the *TYW1*-KO cells and the wild-type ones in neurite arborization. Each experiment was repeated three times.

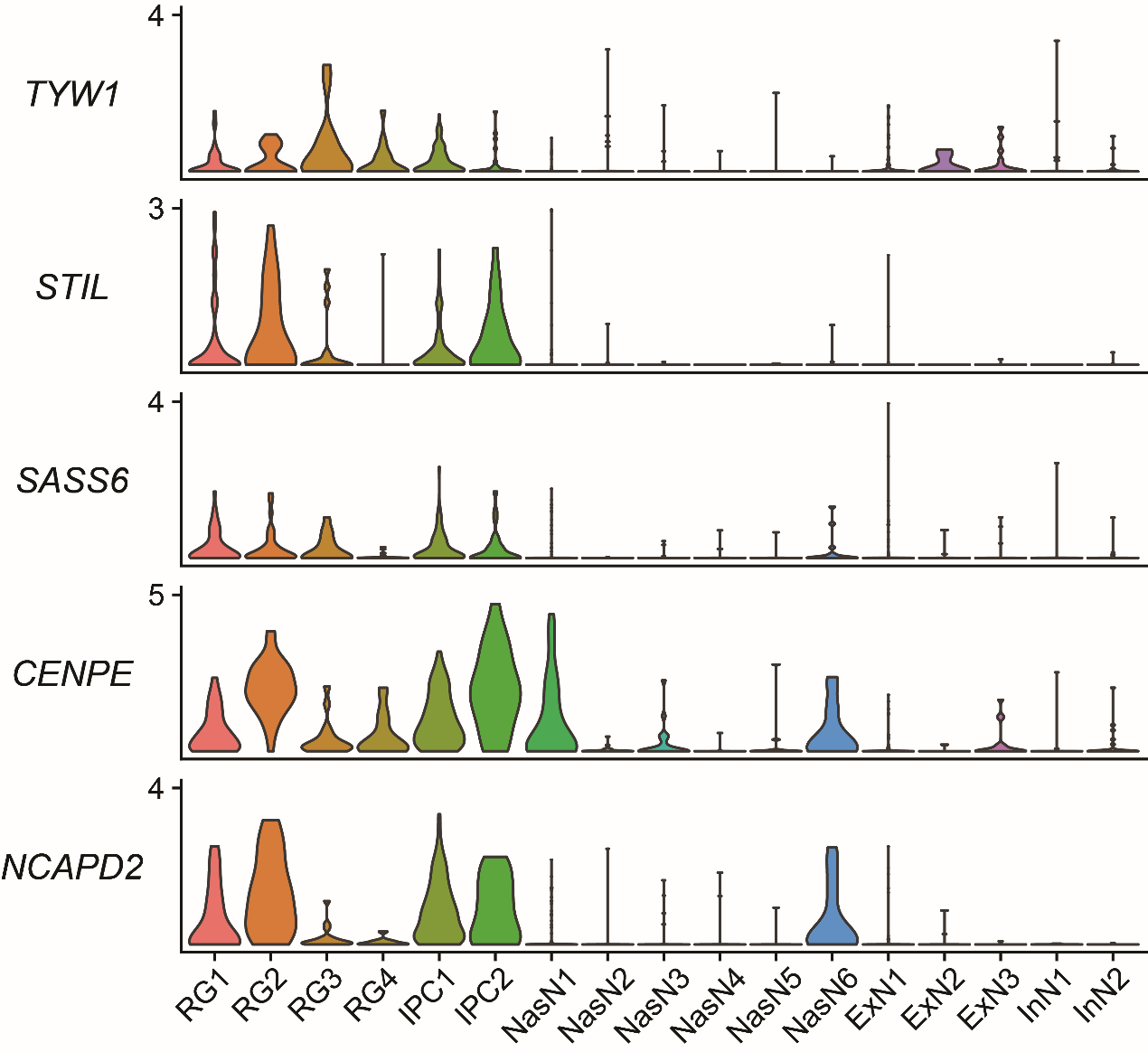

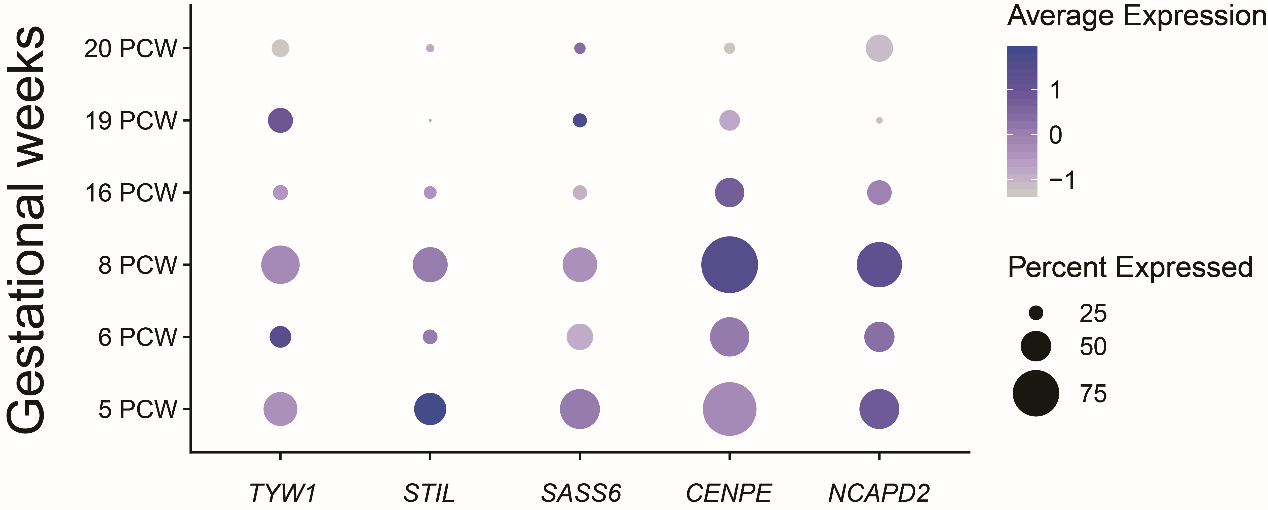

**Supplementary figure 15:** Spatiotemporal mRNA expression patterns of *TYW1*, *STIL*, *SASS6*, *CENPE* and *NCAPD2* were depicted in the different neuronal cell types (above) and gestational weeks (below) ^3^. RG: radial glial cells. IPC: intermediate progenitor cells. NasN: nascent neurons. ExN: excitatory neuron. InN: interneuron. PCW: postconceptional weeks. The R package Seurat (3.1.0) was applied to analyze the single cell data and generate the figures ^4^.

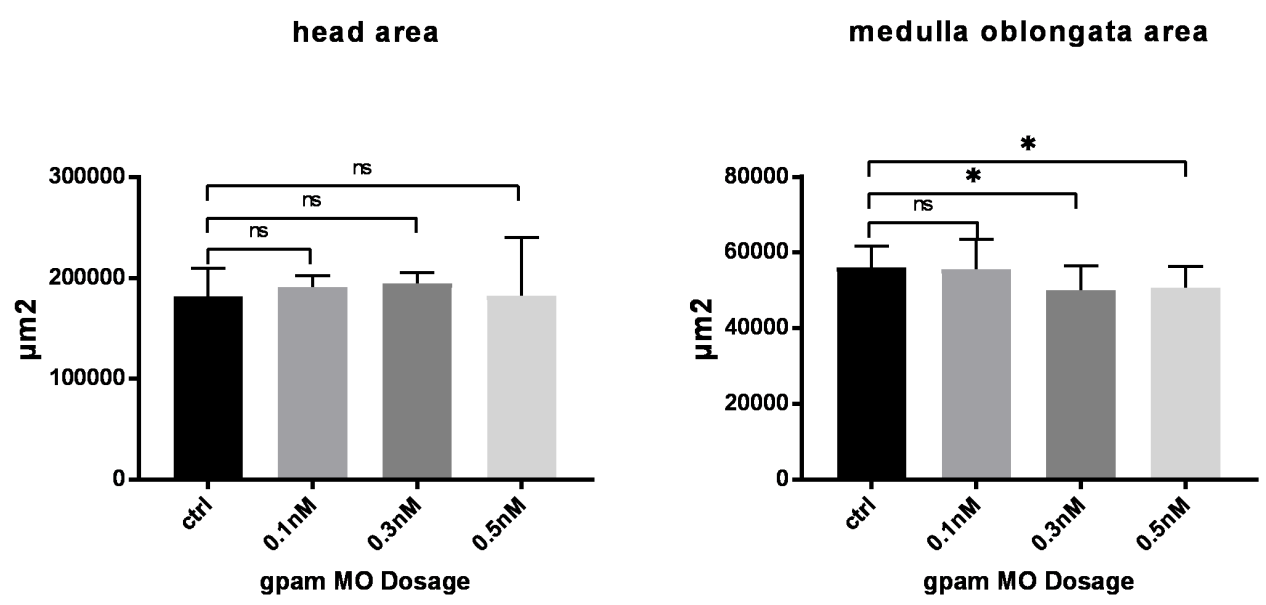

**Supplementary figure 16:** The dosage curves of *gpam*-MO1 knockdown (a splice-blocking Morpholino) efficiency were plotted by measuring the head area and medulla oblongata area of zebrafish larvae at 5 dpf. N = 10. * p < 0.1, unpaired t-test.

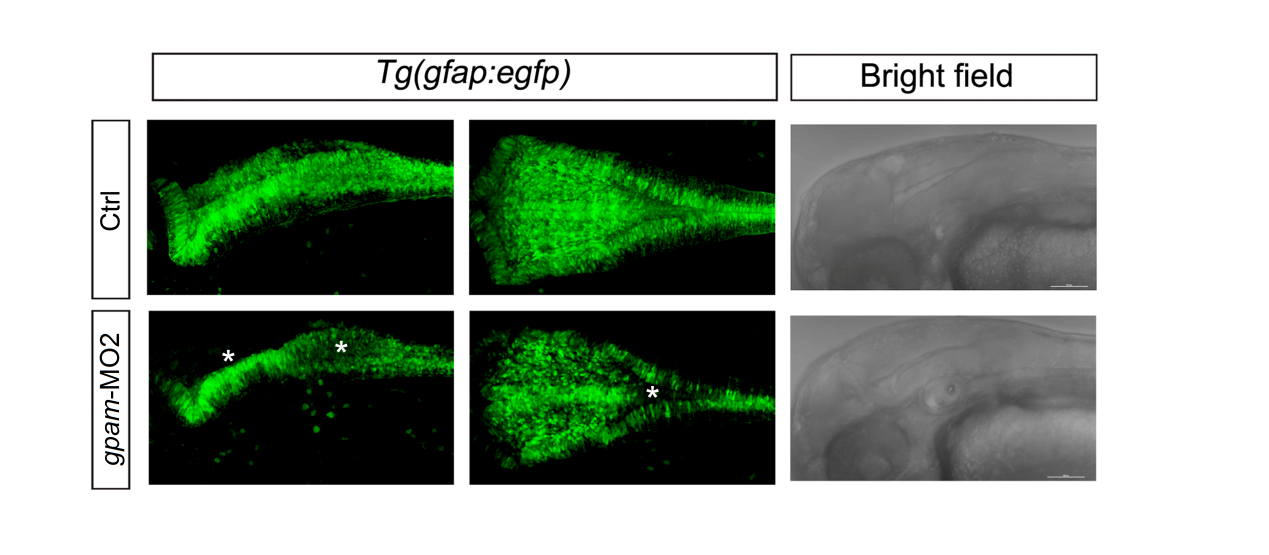

**Supplementary figure 17:** Confocal imaging of egfp^+^ cells in the transgenic zebrafish *Tg(gfap:egfp)* embryos at 48 hpf showed significantly reduced egfp^+^ cells in the *gpam* knockdown group (*gpam*-MO2, a translation blocking morpholino) particularly in the hindbrain (labeled by white stars).
