## Supplementary table 5 Medical charts of the patients with TYW1 and GPAM defects for "In-depth analysis reveals complex molecular etiology of idiopathic cerebral palsy"

**Table 1: Phenotype of patients with compound heterozygous *TYW1* variants.**

| **Characteristics and Symptoms** | **HPO ID ^a^** | **Patient** | **Patient** |
| --- | --- | --- | --- |
| Pedigree ID | N.A. | II.1 | II.2 |
| Origin | N.A. | China | China |
| Gender | N.A. | female | male |
| Age at last assessment (years) | N.A. | 14 | 8.6 |
| **Inheritance** | | | |
| Autosomal Recessive | [0000007](https://hpo.jax.org/app/browse/term/HP:0000007) | + | + |
| **Head & Neck** | | | |
| Microcephaly (head circumference < 2 S.D.) | 0000252 | + | + |
| **Neurologic** | | | |
| Cerebral palsy | [0100021](https://hpo.jax.org/app/browse/term/HP:0100021) | + | + |
| Spastic tetraplegia | 0002510 | + | - |
| Nonprogressive cerebellar ataxia | 0002470 | - | + |
| Athetoid cerebral palsy | 0011445 | + | - |
| Widened cerebral subarachnoid space | 0012766 | N.A. | + |
| Ventriculomegaly | 0002119 | N.A. | + |
| Abnormal cerebellum morphology | 0001317 | N.A. | + |
| Intellectual disability, moderate  (IQ equivalent; years at assessment) | 0002342 | +  (< 50; 13.2) | +  (< 40; 7.5) |
| Delayed speech and language | 0000750 | + | + |
| Delayed fine motor development | 0010862 | + | + |
| Delayed gross motor development | 0002194 | + | + |

**Table 2: Phenotype of patient with compound heterozygous *GPAM* variants.**

| **Characteristics and Symptoms** | **HPO ID ^a^** | **Patient** |
| --- | --- | --- |
| Pedigree ID | N.A. | II.1 |
| Origin | N.A. | China |
| Gender | N.A. | female |
| Age at last assessment (years) | N.A. | 3.6 |
| **Inheritance** | | |
| Autosomal Recessive | [0000007](https://hpo.jax.org/app/browse/term/HP:0000007) | + |
| **Neurologic** | | |
| Cerebral palsy | [0100021](https://hpo.jax.org/app/browse/term/HP:0100021) | + |
| Spastic tetraplegia | 0002510 | + |
| Delayed gross motor development | 0002194 | + |
| Delayed fine motor development | 0010862 | + |
| Focal white matter lesions | 0007042 | + |
| Abnormality of the internal capsule | 0012502 | + |

^a^ Medical records and clinical features are presented according to the recommended schematics and nomenclature of the Clinical Synopsis in OMIM (www.omim.org) and the Human Phenotype Ontology (HPO; [www.human-phenotype-ontology.org](http://www.human-phenotype-ontology.org)).
